# The hGID^GID4^ E3 ubiquitin ligase complex targets ARHGAP11A to regulate cell migration

**DOI:** 10.1101/2023.07.20.549906

**Authors:** Halil Bagci, Martin Winkler, Federico Uliana, Jonathan Boulais, Weaam I Mohamed, Sophia L Park, Jean-François Côté, Matthias Peter

**Author notes:** Address correspondence to: Matthias Peter, Institute of Biochemistry, Department of Biology Otto-Stern-Weg 3, 8093 Zürich Switzerland.

## Abstract

The human CTLH/GID (hGID) complex emerged as an important E3 ligase regulating multiple cellular processes, including cell cycle progression and metabolic activity. However, the range of biological functions controlled by hGID remains unexplored. Here, we show that the hGID substrate receptor GID4 regulates cell growth and migration. Biochemical and cellular assays combined with proximity-dependent biotinylation (BioID2) revealed that the hGID^GID4^ E3-ligase targets the Rho-GAP ARHGAP11A for degradation. Depletion of GID4 or impeding the GID4 substrate binding pocket impairs motility and directed cell movement, whereas knockdown of ARHGAP11A significantly restores the cell migration defect. We found that GID4 controls cell migration by degrading ARHGAP11A thereby preventing its accumulation at the cell periphery where it inactivates RhoA activity. Together, we identified a unique function for GID4, as well as a wide range of substrate profiles beyond Pro/N-degron motifs, which pave the way for deciphering additional pathways regulated by hGID E3 ligase activity through its GID4 substrate receptor.

## Introduction

The ubiquitin-proteasome system (UPS) is a key protein degradation machinery in eukaryotic cells. Conjugation of ubiquitin to target proteins is achieved by three coordinated enzymatic reactions, governed by an activating E1, conjugating E2, and ligating E3 enzymes. E3 ligases perform the critical function of substrate recognition, in some cases by detecting specific short amino acid motifs called degrons. Ubiquitin conjugation to substrate proteins regulates various cellular processes, including cellular homeostasis, metabolism and cell cycle progression (Brandon Croft, 2015). Dysfunctions in the UPS, including mutations in the ubiquitin machinery or in substrate recognition motifs, have been associated with a broad spectrum of pathological conditions including cancer and metabolic diseases (Kitamura, 2023).

In yeast, the UPS tightly controls the metabolic switch from gluconeogenesis to glycolysis. This process involves Glucose-induced degradation deficient (GID) proteins, also known as the C-terminal to LisH (CTLH), which form a multi-subunit RING-domain-containing E3 ligase (Santt *et al*, 2008). Biochemical and structural analysis revealed that the yeast GID complex is composed of seven subunits, and four such units assemble into a stable tetramer. The catalytic center is formed by the two RING-domain-containing proteins Gid2 and Gid9, which are held together by the scaffold Gid8. Gid5 recruits different substrate-receptors including Gid4, Gid10 and Gid11. Gid4 promotes proteasomal degradation of excess gluconeogenic enzymes such as fructose-1,6-biphosphate 1 (Fbp1) or malate dehydrogenase (Mdh2) (Chen *et al*, 2017; Dong *et al*, 2020). These substrates are recognized via a Pro/N-end degron motif, which docks into a conserved Gid4 binding pocket. Similarly, Gid10 and Gid11 target distinct sets of substrates that regulate specific metabolic transitions (Langlois *et al*, 2022; Kong *et al*, 2021).

The GID/CTLH E3 ligase complex is evolutionary conserved, and all yeast subunits have closely related counterparts in higher eukaryotes (Salemi *et al*, 2017; Lampert *et al*, 2018; Maitland *et al*, 2019). RanBP9 (Gid1), RMND5A (Gid2), ARMC8 (Gid5), TWA1 (Gid8) and MAEA (Gid9) are ubiquitously expressed and assemble into multimeric complexes localizing to the nucleus and cytoplasm (Kobayashi *et al*, 2007). The two RING-domain containing subunits RMND5A and MAEA linked by TWA1 form the catalytic trimer (Lampert *et al*, 2018), which assemble with other subunits such as WDR26 (Gid7), RanBP9/RanBP10 (Gid1), MKLN1, GID4 (Gid4), ARMC8 (Gid5) and YPEL5 (Fig 1A) (Kobayashi *et al*, 2007; Lampert *et al*, 2018). While structural studies revealed important insight into the mechanism and assembly of hGID E3 ligase complexes (Sherpa *et al*, 2021; Mohamed *et al*, 2021), its physiological functions and substrates remain poorly characterized.

**Fig 1.**
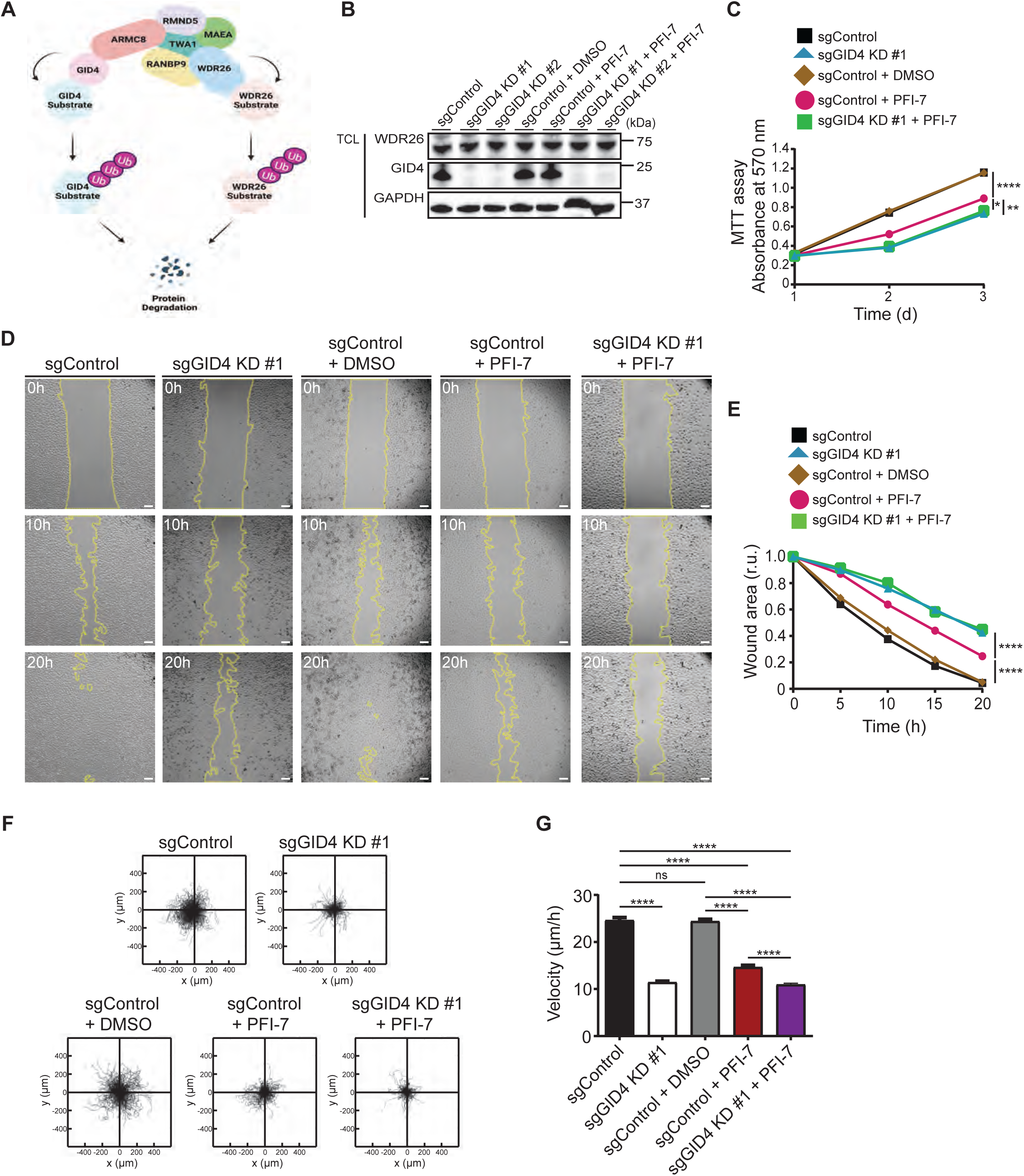
Loss of GID4 leads to cell migration defect. A Schematic of the hGID E3 ligase complex. The hGID complex includes multiple subunits and comprises two distinct substrate receptors (GID4 and WDR26), two RING E3 ligases (RMND5 and MAEA), and other subunits including TWA1, ARMC8 and RANBP9. Each substrate receptor can target a specific set of substrates for protein degradation via ubiquitination. Figure created with BioRender.com. B Western blots of HeLa total cell lysates (TCL) showing GID4 and WDR26 protein expression. Lysates were prepared from a stable clone without sgRNA (sgControl), a stable clone with a pool of four sgRNAs targeting *Gid4* (sgGID4 KD #1), a second stable clone with a pool of four sgRNAs targeting *Gid4* (sgGID4 KD #2), sgControl treated with DMSO (10 µM), sgControl treated with PFI-7 (10 µM), sgGID4 KD #1 treated with PFI-7 (10 µM) and sgGID4 KD #2 treated with DMSO (10 µM) or PFI-7 (10 µM). Blots were probed where indicated with antibodies to GID4, WDR26 and GAPDH as a loading control. Blot shown is representative of three independent experiments. C MTT assay in HeLa cells measuring absorbance at 570nm to indicate cell metabolic activity during 1, 2 or 3 days (d) for lysates derived as in Fig 1B. sgControl or sgGID4 KD #1 cells were either untreated or treated with DMSO (10 µM) or PFI-7 (10 µM). Data values at day 3 were analyzed for statistical significance and shown as mean ±SD (n=3 independent experiments; 3 biological replicates were performed for each experiment). The indicated *P*-values were calculated by one-way ANOVA, followed by Bonferroni’s multiple comparisons test. \**P*≤0.05, \*\**P*≤0.01, \*\*\*\**P*≤0.0001 D Representative brightfield images acquired over time (h) of a wound healing assay with HeLa sgControl or sgGID4 KD #1 cells, either untreated or treated with DMSO (10 µM) or PFI-7 (10 µM). Cells were grown to a monolayer with a defined cell-free gap established by silicone insert. The silicone insert was removed and images were acquired at 1 h intervals from initial time (0 h) to end (20 h). Images acquired at time 0 h, 10 h and 20 h are shown. The wound area was selected using the freehand selection tool (ImageJ) and is outlined in yellow. Scale bars, 100 µm. E Line graph showing wound area measurements in relative units (r.u.) over time (h) from images acquired as in Fig 1D with wound area selected as in Fig 1D and normalized to starting cell-free wound area at time 0h. Data values at 20 hours were analyzed for statistical significance and are shown as mean ±SD (n=3 independent experiments; 4 measurements were performed for each wounded area). The indicated *P*-values were calculated by one-way ANOVA, followed by Bonferroni’s multiple comparisons test. \*\*\*\**P*≤0.0001 F Plots showing a 24 h period of merged individual RPE1 cell trajectories set to a common origin at the intersection of y (µm) and x (µm) axes for sgControl and sgGID4 KD #1 cells, either untreated or treated with DMSO (10 µM) or PFI-7 (10 µM). Images were acquired at 30 min intervals for 24 h, analyzed using a manual tracking plugin and chemotaxis tool (Ibidi) in ImageJ software. G Bar graph showing cell velocity (µm/h) of RPE1 cells from data acquired and analyzed as in Fig 1F. Data values are shown as mean ±SD (n=3 independent experiments; 200 cells were analyzed for each condition). The indicated *P*-values were calculated by one-way ANOVA, followed by Bonferroni’s multiple comparisons test. ns (not significant), \*\*\*\**P*≤0.0001

Comprehensive phage display screens and peptide binding assays demonstrated that human GID4 (hGID4) subunit binds a variety of short motifs via a conserved pocket (Chrustowicz *et al*, 2022). A chemical compound, PFI-7, blocks this binding pocket, thereby preventing hGID4 interaction with N/Pro-degron containing targets, such as DNA-helicases DDX21 or DDX50 (Owens *et al*, 2023). Nevertheless, it remains unclear whether hGID4 primarily degrades Pro/N-end containing substrates *in vivo*, as the GID4 substrate ZMYND19 lacks an N-terminal degron compatible with the proposed consensus motif (Mohamed *et al*, 2021). Moreover, GID4 may not be the only substrate-receptor of the hGID E3 ligase, as depletion of WDR26/Gid7 but not hGID4 stabilizes the tumor suppressor Hbp1 (Fig 1A) (Lampert *et al*, 2018; Mohamed *et al*, 2021).

Although the biological functions of the hGID E3 ligase are only beginning to emerge, its activity has been implicated in regulating cell proliferation, metabolism, embryonic development and cell differentiation. Moreover, several hGID subunits such as RanBP9, MKLN1 and WDR26 have been associated with cell migration and adhesion (Maitland *et al*, 2022), with altered expression levels in many invasive and metastatic cancer cells (Ye *et al*, 2016). Cell migration is a highly integrated multistep process driven by spatio-temporal control of membrane protrusions and actin polymerization at the leading edge of the cell. Subsequent steps include adhesion to matrix contacts, contraction of the cytoplasm, release from contact sites and recycling of membrane receptors from the rear to the front of the cell. Actin dynamics are regulated by the activity of Rho GTPases through the opposing actions of a large family of guanine nucleotide exchange factors (GEFs) and GTPase-activating proteins (GAPs). Interestingly, RanBP9 interacts with ß-integrins and promotes cell attachment and spreading (Woo *et al*, 2012), while MKLN1 and WDR26 regulate the activity of Rho-type GTPases (Tripathi *et al*, 2015) (Hasegawa *et al*, 2020). However, how hGID-complexes regulate Rho-type GTPases and influence cell migration and invasion remains unclear.

Here we employed an integrative approach combining cellular phenotyping and systematic BioID2-based mass spectrometry to uncover physiological hGID substrates involved in cell growth and migration. Interestingly, our findings demonstrate that GID4 tightly controls cell migration by regulating RhoA activity, which is achieved through ubiquitination and subsequent degradation of the RhoGAP ARHGAP11A. Indeed, abrogation of GID4 expression or inhibition of its substrate binding pocket leads to the accumulation of ARHGAP11A at the cell periphery, where it colocalizes with F-actin protrusions and inactivates RhoA. Collectively, our study represents a valuable resource that recapitulates the transient interactome of GID4, altered by proteasomal degradation and its substrate-binding pocket. Among the interactors, we validated the relationship between GID4 and ARHGAP11A, which functions as a physiological substrate of the hGID^GID4^ E3 ligase regulating cell growth and migration.

## Results

### GID4 is required for efficient cell growth and migration

To investigate the role of the hGID E3-ligase and in particular its substrate receptor GID4 in regulating cell growth and proliferation, we generated stable, doxycycline (DOX)-inducible GID4 knockdown (KD) HeLa and RPE1 cell lines using the CRISPR-Bac system (Schertzer *et al*, 2019). Briefly, HeLa or RPE1 cells were transfected with a pool of four single guide RNAs (sgRNAs) targeting the *GID4* gene or without a sgRNA for control (sgControl). Stable integration of the sgRNA and DOX-inducible PB_tre_Cas9 vector was selected using G418 and hygromycin for HeLa, or G418 and puromycin for RPE1 cells. Independent clones were expanded and efficient GID4 knockdown was confirmed by immunoblotting after DOX-induction for 96 hours. For both the HeLa and RPE1 cell lines, two validated clones termed sgGID4 KD #1 and sgGID4 KD #2 were further characterized and used throughout this study. Importantly, while GID4 was efficiently depleted, WDR26 levels remained unchanged for both the HeLa and RPE1 cell lines, confirming specificity of the sgGID4 and stability of the remaining hGID complex (Figs 1B and EV1A). We also used the recently described GID4-inhibitor PFI-7, which binds to a structurally defined GID4 pocket, thereby blocking access of N-terminal degron motifs (Owens *et al*, 2023). Compared to DMSO controls, addition of PFI-7 to sgControl HeLa or RPE-1 cells did not alter GID4 or WDR26 levels, respectively (Figs 1B and EV1A).

To uncover cellular functions of the hGID^GID4^ complex, we first tested if GID4 depletion affects proliferation of Hela cells. Interestingly, cell lines lacking GID4 or treated with the GID4-inhibitor PFI-7 showed approximately two-fold reduced growth rates compared to control cells or DMSO alone, as measured by MTT-absorbance at 570nm (Figs 1C and EV1B). Addition of the PFI-7 compound to GID4-depleted cell lines did not further enhance this proliferation defect, confirming that PFI-7 is specific and GID4 is the relevant PFI-7 target underlying this phenotype.

To confirm and extend these results, we performed wound healing assays to examine GID4 function in directed cell migration. We observed more than six-fold delay of both sgGID4 KD #1 and sgGID4 KD #2 HeLa cells to close the cell-free area compared to sgControl (Figs 1D and E and EV1C and D). Similarly, the wound area of PFI-7-treated sgControl HeLa cells was more than four-fold larger than sgControl cells treated with DMSO (Fig 1D). The wound healing response of PFI-7-treated cells was less pronounced compared to both sgGID4 KD cell lines, while PFI-7 addition to GID4-depleted cells did not enhance the phenotype. These results suggest that blocking the GID4 substrate binding pocket is less effective as compared to cells lacking GID4.

Delayed wound closure could be a consequence of reduced proliferation or a combination of reduced proliferation and impaired cell migration. To investigate whether decreased cell migration may contribute to the wound healing defect, we carried out single cell tracking assays using RPE1 cells. Interestingly, we observed that GID4-depleted RPE1 cells exhibited more than two-fold decreased velocity compared to sgControl cells (Figs 1F and G and EV1E-G). Likewise, PFI-7-treated sgControl cells also showed reduced velocity compared to DMSO-treated sgControls, albeit less pronounced than observed with untreated or PFI-7-treated sgGID4 KD cells. Taken together, we conclude that GID4 is required for efficient cell motility, suggesting the hGID^GID4^ E3 ligase complex regulates targets specifically involved in this process.

### BioID2-mediated proximity labeling identifies potential GID4 substrates

To decipher the GID4 proximal protein interaction network and explore potential GID4 substrates regulating cell migration as well as other biological functions, we carried out a proximity-dependent biotinylation screen (Fig 2A). To achieve this goal, we first generated stable Flp-In T-REx HeLa cells expressing a BirA2-Flag-GID4 fusion protein (BioID2-GID4) in a tetracycline-inducible manner. As a control, we generated Flp-In T-REx HeLa cell lines expressing BirA2-Flag-EGFP (BioID2-GFP) or an empty vector. Moreover, to distinguish potential substrates from general interactors/regulators, we generated a BirA2-Flag-GID4^E237A^ fusion cell line (BioID2-GID4^E237A^), which harbors a specific point mutation known to abolish substrate binding (Dong *et al*, 2018). Immunoblotting with FLAG-antibodies confirmed that the BioID2-GID4 or BioID2-GID4^E237A^ proteins are expressed at comparable levels, and they stably assemble into the hGID complex measured by co-immunoprecipitation with its catalytic subunit MAEA (Figs EV2A and B). Finally, treating the BioID2-GID4 and BioID2-GID4^E237A^ cell lines with tetracycline and biotin demonstrated that both fusion proteins efficiently trigger biotinylation of endogenous proteins in their vicinity (Fig EV2A).

**Fig 2.**
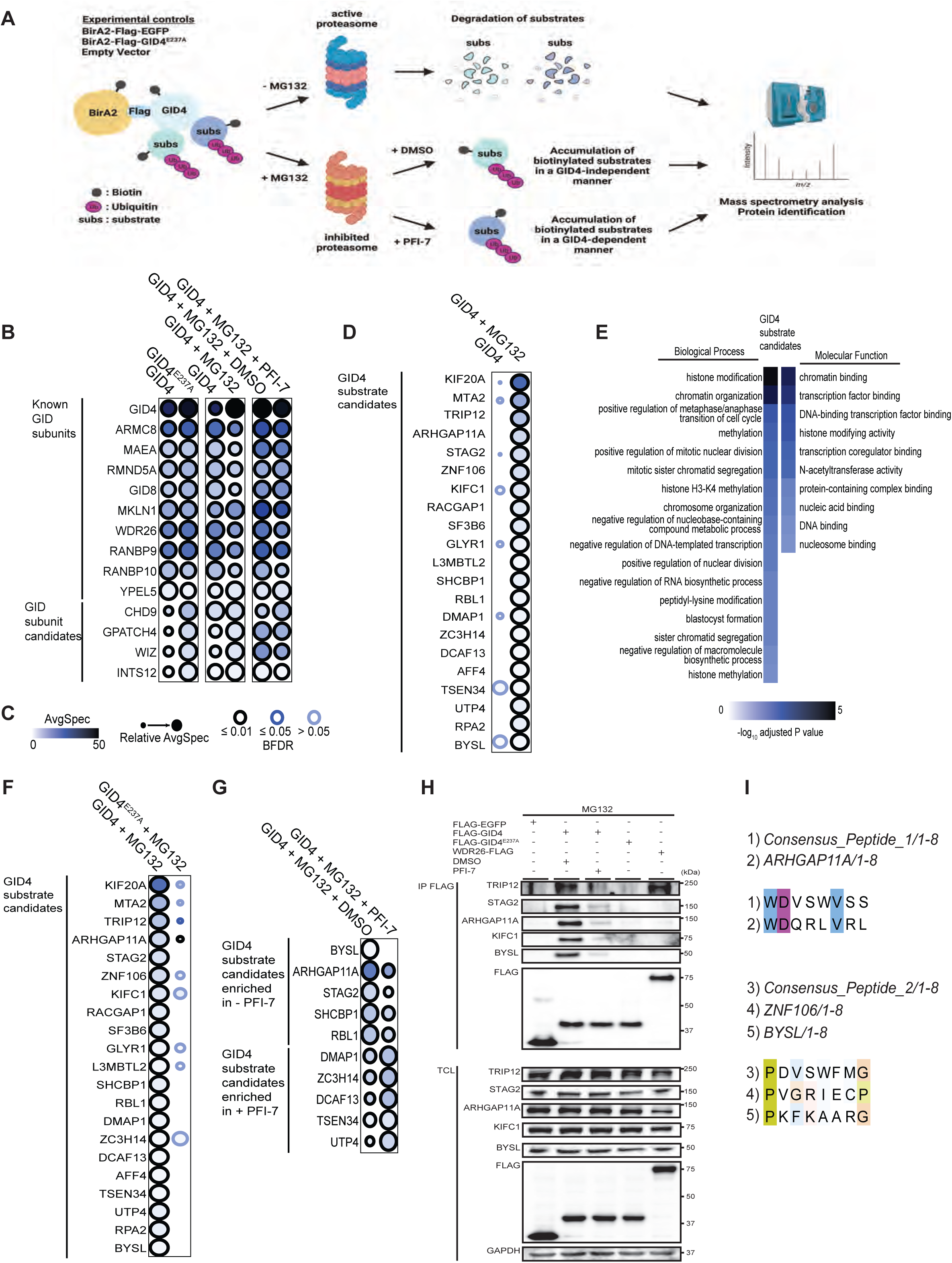
Proximity labeling by BioID2 identifies potential GID4 substrates. A Workflow of the BioID2 pipeline to identify GID4 substrates. Flp-In T-REx HeLa cell lines expressing either BirA2-Flag-GID4 (BioID2-GID4), BirA2-Flag-GID4^E237A^ (BioID2-GID4^E237A^), BirA2-Flag-EGFP (BioID2-GFP), or empty vector control BirA2-Flag alone (Empty Vector) as bait proteins were treated with tetracycline (1 µg/ml) and biotin (50 µM) in the presence or absence of MG132 (5 mM), with or without DMSO (10 µM) or PFI-7 (10 µM) for 24 hours. Cell lysates were incubated with streptavidin beads, and bead-associated proteins were digested by trypsin. The resulting peptides were analyzed by mass spectrometry (MS) for protein identification. Schematic adapted from (Gingras *et al*, 2019). Figure created with BioRender.com. B Dot plots of BioID2 interacting proteins identified as in Fig 2A displaying analysis and quantification (ProHits) of output MS data (SAINT) as indicated in Fig 2C. HeLa BioID2-GID4 or BioID2-GID4^E237A^ cell lines expressing the respective BirA2-Flag-tagged bait protein (GID4, GID4^E237A^) were either untreated, or treated with MG132 (5 mM), and/or DMSO (10 µM) or PFI-7 (10 µM). C Legend for dot plot data in (Figs 2B and D-G) displaying analysis and quantification (ProHits) of output MS data (SAINT). The average spectral counts are represented by the node color. The edge color shows the confidence score of the BioID2 interaction (BFDR ≤ 1% considered as high confidence, 1% < BFDR ≤ 5% as medium confidence or BFDR > 5% as low confidence). The relative abundance of prey is depicted by the circle size according to the biggest node size and proportionally scaled for the other preys. D Dot plots of BioID2 interacting proteins identified as in Fig 2A displaying analysis and quantification (ProHits) of output MS data (SAINT) as indicated in Fig 2C. HeLa BioID2-GID4 cell lines expressing BirA2-Flag-tagged bait protein (GID4) was either untreated or treated with MG132 (5 mM). E Heatmap of gene ontology (GO) term analysis for GID4 substrate candidates representing the enrichment score for selected biological processes and molecular functions (top) as imported from g:Profiler. The enrichment score for each GO term is shown as the –log10 of adjusted *P* values, displayed by the legend with varying color intensities (bottom). F Dot plots of BioID2 interacting proteins identified as in Fig 2A displaying analysis and quantification (ProHits) of output MS data (SAINT) as indicated in Fig 2C. HeLa BioID2-GID4 or BioID2-GID4^E237A^ cell lines expressing the respective BirA2-Flag-tagged bait protein (GID4, GID4^E237A^) were treated with MG132 (5 mM). G Dot plots of BioID2 interacting proteins identified as in Fig 2A displaying analysis and quantification (ProHits) of output MS data (SAINT) as indicated in Fig 2C. HeLa BioID2-GID4 cells lines expressing BirA2-Flag-tagged bait protein (GID4) was treated with MG132 (5 mM) and either DMSO (10 µM) or PFI-7 (10 µM). H Western blots of bound co-immunoprecipitated proteins (IP-FLAG) and unbound total cell lysate (TCL) from Flp-In T-Rex HeLa cell lines expressing FLAG-EGFP, FLAG-GID4, FLAG-GID4^E237A^ or WDR26-FLAG as bait proteins. Where indicated, samples were either untreated, or treated with DMSO (10 µM), PFI-7 (10 µM) or MG132 (5 mM). Blots were probed where indicated with antibodies to FLAG, or to endogenous TRIP12, STAG2, ARHGAP11A, KIFC1 or BYSL, and GAPDH as a loading control. I Sequence alignment showing the first 8 amino acids at the N-terminus of consensus motifs from GID4-binding peptides (Consensus motifs from (Chrustowicz *et al*, 2022)) with ARHGAP11A or ZNF106 and BYLS.

To identify proximal candidate substrates of GID4, we next incubated the cell lines with tetracycline and biotin for 24 hours and affinity-isolated biotinylated proteins from cell extracts using Streptavidin beads. After elution, biotinylated proteins were digested on beads, identified by mass-spectrometry (MS), and quantified by spectral counts. We further used the SAINT algorithm (Choi *et al*, 2011) comparing intensity of proteins at different conditions versus experimental controls. To distinguish substrates from other hGID4-interacting proteins, we performed the BioID assays in cells treated or not with the proteasome inhibitor MG132 (Fig 2A). In addition, we co-treated cells with MG132 and the PFI-7 inhibitor to classify proteins that interact with GID4 in a substrate-binding-pocket dependent manner. We successfully recovered multiple known hGID subunits with both BioID2-GID4 and BioID2-GID4^E237A^ baits (Figs 2B and C) in the absence or presence of MG132 or PFI-7, confirming that GID4 engages with the hGID complex independently of proteasome function or substrate binding. Interestingly, we detected several additional interactors with a high-confidence score in all conditions (BFDR < 0.01), including CHD9, GPATCH4, WIZ or INTS12, suggesting their potential role as hGID subunits and/or regulators.

Next, we extended our search for proteins that exhibit substrate behavior. We first identified GID4 interacting proteins that are significantly enriched in MG132-treated BioID2-GID4 cells. We identified multiple proteins with high spectral counts and confidence scores (BFDR < 0.01) including KIF20A, MTA2, TRIP12, ARHGAP11A, STAG2, ZNF106, KIFC1, RACGAP1 (Figs 2D and C). Gene ontology (GO) analysis of the interactors enriched in the MG132-treated BioID2-GID4 cells revealed that most of these putative GID4 substrates are involved in chromatin organization, gene expression, histone modification and chromosome segregation, suggesting a role in chromatin regulation and cell division (Fig 2E), and implicating a nuclear role of GID4 in cells. Importantly, most of the GID4 substrate candidates require an intact GID4-substrate binding pocket as their spectral counts are greatly reduced in BioID2-GID4^E237A^ expressing cell lines (Figs 2F and C). To corroborate these data, we performed another BioID2 screen in MG132-treated BioID2-GID4 cells with or without PFI-7 (Figs 2G and C). Interestingly, we found candidates that were either reduced, enriched or unchanged upon PFI-7 treatment. Among the ones with reduced spectral counts were BYSL, ARHGAP11A, and STAG2, confirming that they may bind to GID4 via its substrate binding pocket (Figs 2G and C). Surprisingly, many candidates that lost their interaction to the BioID2-GID4^E237A^ mutant were unaffected by PFI-7 treatment (Table EV1), suggesting that blocking substrate binding by PFI-7 is not complete, or alternatively, that the E237A mutant may disrupt more than the GID4 substrate pocket.

To further validate GID4 candidate substrates identified by BioID2 screening, we immunoprecipitated FLAG-tagged wild-type GID4 or its E237A mutant and probed for co-immunoprecipitation of selected candidates using specific antibodies (Fig 2H). We also tested whether their binding is altered by the PFI-7 inhibitor, and included FLAG-tagged WDR26 to confirm that these substrates engage the hGID complex via GID4 and not the alternate WDR26 substrate receptor. In line with our BioID2 results, BYSL, KIFC1, ARHGAP11A and STAG2 readily co-immunoprecipitated with GID4, and their interaction with the GID4-E237A mutant was significantly diminished. While these proteins specifically co-immunoprecipitated with GID4 but not WDR26, TRIP12 interacted with both substrate receptors, suggesting that it might be a GID4 substrate that also binds WDR26 or may exclusively interact with oligomeric hGID assemblies containing both subunits.

Protein sequence alignment of the GID4 substrate candidates showed that ARHGAP11A encompasses an N-terminal motif (WDQRLVRL) matching the peptide binding consensus determined previously (Chrustowicz *et al*, 2022), implying that this degron may target ARHGAP11A to the GID4 binding pocket (Fig 2I). Likewise, ZNF106 and BYSL conform to an alternative Pro/N-degron consensus motif, while other putative substrates do not fit the available criteria, even though their binding seems dependent on an intact GID4 binding pocket.

Taken together, this comprehensive BioID2 analysis identified numerous GID4 interactors that (1) bind as predicted for hGID subunits and/or regulators, or (2) exhibit substrate-like behavior, where their interaction is increased in the presence of MG132 and altered by an intact GID4 substrate-pocket.

### ARHGAP11A is ubiquitinated and degraded by a GID4-dependent mechanism

We next used the STRING database (Szklarczyk *et al*, 2019) to highlight relevant protein complexes or clusters among the GID4 substrate candidates that may be functionally important to explain the role of GID4 in cell proliferation and cell migration. Interestingly, this analysis revealed that ARHGAP11A, KIFC1, RACGAP1, KIF20A and SHCBP1 comprise an interrelated cluster regulating cytoskeletal dynamics and cytokinesis as well as microtubule binding and motor activity (Fig 3A). We thus tested whether these GID4 substrate candidates are degraded *in vivo* in a GID4-dependent manner. To this end, we treated HeLa sgGID4 KD or sgControl cell lines with the translation-inhibitor cycloheximide (CHX), and assessed the half-life of ARHGAP11A, KIFC1 and RACGAP1 by immunoblotting (Figs 3B and C). Interestingly, while all three proteins are degraded with a half-life below 4 hours, only ARHGAP11A was stabilized in the absence of GID4 (Figs 3B and C). To corroborate these data, we analyzed ARHGAP11A and KIFC1 levels in CHX-treated HeLa sgGID4 KD or sgControl cell lines that were also treated with PFI-7 or DMSO. Indeed, ARHGAP11A but not KIFC1 was stabilized in the presence of PFI-7, implying that binding to GID4 is required for ARHGAP11A degradation *in vivo* (Figs 3D and E). In contrast, KIFC1 and RACGAP1 may indirectly binding to GID4, as their degradation is mediated by a GID4-independent mechanism.

**Fig 3.**
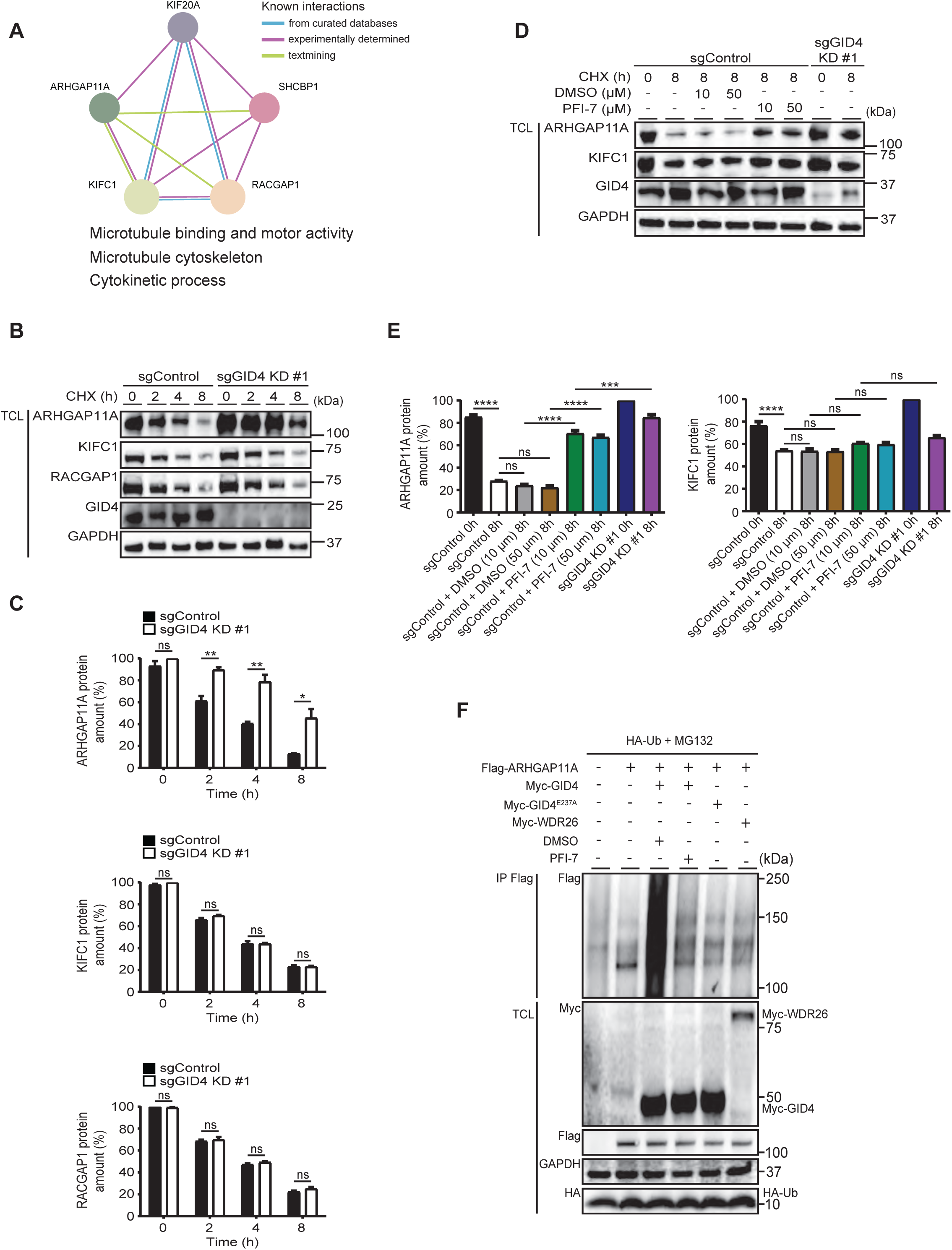
ARHGAP11A acts as a GID4 degradation substrate. A Protein network representation of the five selected high-confidence, MG132-enriched interactors of GID4. Molecular functions or biological processes related to the network are imported from g:Profiler and listed below the network. The –log10 of adjusted *P* values for microtubule binding, microtubule motor activity or cytokinetic process are 2.306, 1.650 and 1.522, respectively. Protein network was generated with MCL clustering using STRING v11.5. B Western blots of a Cycloheximide (CHX) chase assay with total cell lysates (TCL) of sgControl or sgGID4 KD #1 HeLa cells. CHX (20 µg/ml) was added at the indicated time points (0 h – 8 h). Blots were probed where indicated with antibodies to endogenous GID4, ARHGAP11A, KIFC1 or RACGAP1, and GAPDH as a loading control. C Bar graphs of a CHX chase assay prepared as in Fig 3B, showing the amount (%) of ARHGAP11A (top), KIFC1 (middle) or RACGAP1 (bottom) protein over time (h), normalized to starting protein amount at time 0h. Data values are shown as mean ±SEM (n=3 independent experiments). The indicated *P*-values were calculated by a two-tailed Student’s *t* test. ns (not significant), \**P*≤0.05, \*\**P*≤0.01 D Western blots of a Cycloheximide (CHX) chase assay in which total cell lysates (TCL) of sgControl or sgGID4 KD #1 HeLa cells were treated with DMSO or PFI-7 at the indicated dose (μM). CHX (20 µg/ml) was added at the indicated time points (0 h – 8 h). Blots were probed where indicated with antibodies to endogenous GID4, ARHGAP11A, or KIFC1, and GAPDH as a loading control. E Bar graphs of a CHX chase assay prepared as in Fig 3D, showing the amount (%) of ARHGAP11A (left) or KIFC1 (right) protein over time (h), normalized to starting protein amount at time 0h. Data values are shown as mean ±SD (n=3 independent experiments). The indicated *P*-values were calculated by one-way ANOVA, followed by Bonferroni’s multiple comparisons test. ns (not significant), \*\*\**P*≤0.001, \*\*\*\**P*≤0.0001 F Western blots of bound co-immunoprecipitated proteins (IP-FLAG) and unbound total cell lysate (TCL) from HeLa cells expressing Flag-ARHGAP11A and either Myc-GID4, Myc-GID4 ^E237A^, or Myc-WDR26 in the presence of HA-ubiquitin (HA-Ub) and MG132 (5 mM). Where indicated, cells were untreated or treated with DMSO (10 µM) or PFI-7 (10 µM). Blot of bound fraction (IP-FLAG) was probed with anti-Flag antibody. Blot of TCL was probed as indicated with antibodies to Myc-, Flag-, and HA-tags, and with antibody to endogenous GAPDH as a loading control.

To validate that ARHGAP11A is ubiquitinated by hGID^GID4^, we immunoprecipitated FLAG-tagged ARHGAP11A from MG132-treated HeLa cells overexpressing HA-tagged ubiquitin and either Myc-tagged GID4^E237A^ or wild-type GID4 treated with DMSO or PFI-7 to block substrate binding (Fig 3F). For further control, we overexpressed Myc-tagged WDR26. Indeed, immunoprecipitation of FLAG-tagged ARHGAP11A revealed a smear of high-molecular weight species, consistent with ubiquitinated ARHGAP11A (Fig 3F). Importantly, ARHGAP11A polyubiquitination was inhibited by the addition of PFI-7 and the GID4-E237A mutation, and no ubiquitination of ARHGAP11A was observed in cells overexpressing Myc-tagged WDR26. Taken together, these results demonstrate that the hGID^GID4^ E3-ligase ubiquitinates ARHGAP11A, which targets it for rapid degradation by the 26S proteasome.

### GID4-dependent degradation of ARHGAP11A regulates cell migration through RhoA activity

To interrogate whether GID4 regulates cell migration by controlling ARHGAP11A stability, we first abolished the ARHGAP11A expression by siRNA in sgControl HeLa or sgControl RPE1 cells (Figs 4A and B and EV3A and B). Consistent with previous results (Dai *et al*, 2018; Lawson *et al*, 2016), siARHGAP11A-depleted sgControl HeLa or RPE1 cells displayed a profound migration defect, as determined by wound healing assays and single cell velocity measurements (Figs 4C-E). Importantly, these results imply that reduced and increased ARHGAP11A levels, as observed upon loss of GID4 function, both cause defects in cell migration and motility, characteristic of altered GTPase dynamics. To test whether GID4 functionally antagonizes ARGAP11A levels *in vivo*, we treated siARHGAP11A-transfected sgControl HeLa or RPE1 cells with the GID4-inhibitor PFI-7 to reduce ARHGAP11A turnover. Indeed, PFI-7 increased ARHGAP11A protein levels more than two-fold compared to the untreated siARHGAP11A-transfected cells (Figs 4F and G and EV3C and D). Importantly, the partially restored ARHGAP11A levels was sufficient to improve the wound healing and migration velocity defects observed in ARHGAP11A-RNAi-depleted sgControl cells, confirming that GID4 antagonizes ARHGAP11A turnover and function *in vivo* (Figs 4H and I and EV3E-G). Together, these data demonstrate that ARHGAP11A levels are regulated by the hGID^GID4^ E3 ligase, and either loss or gain of ARHGAP11A function leads to cell migration defects.

**Fig 4.**
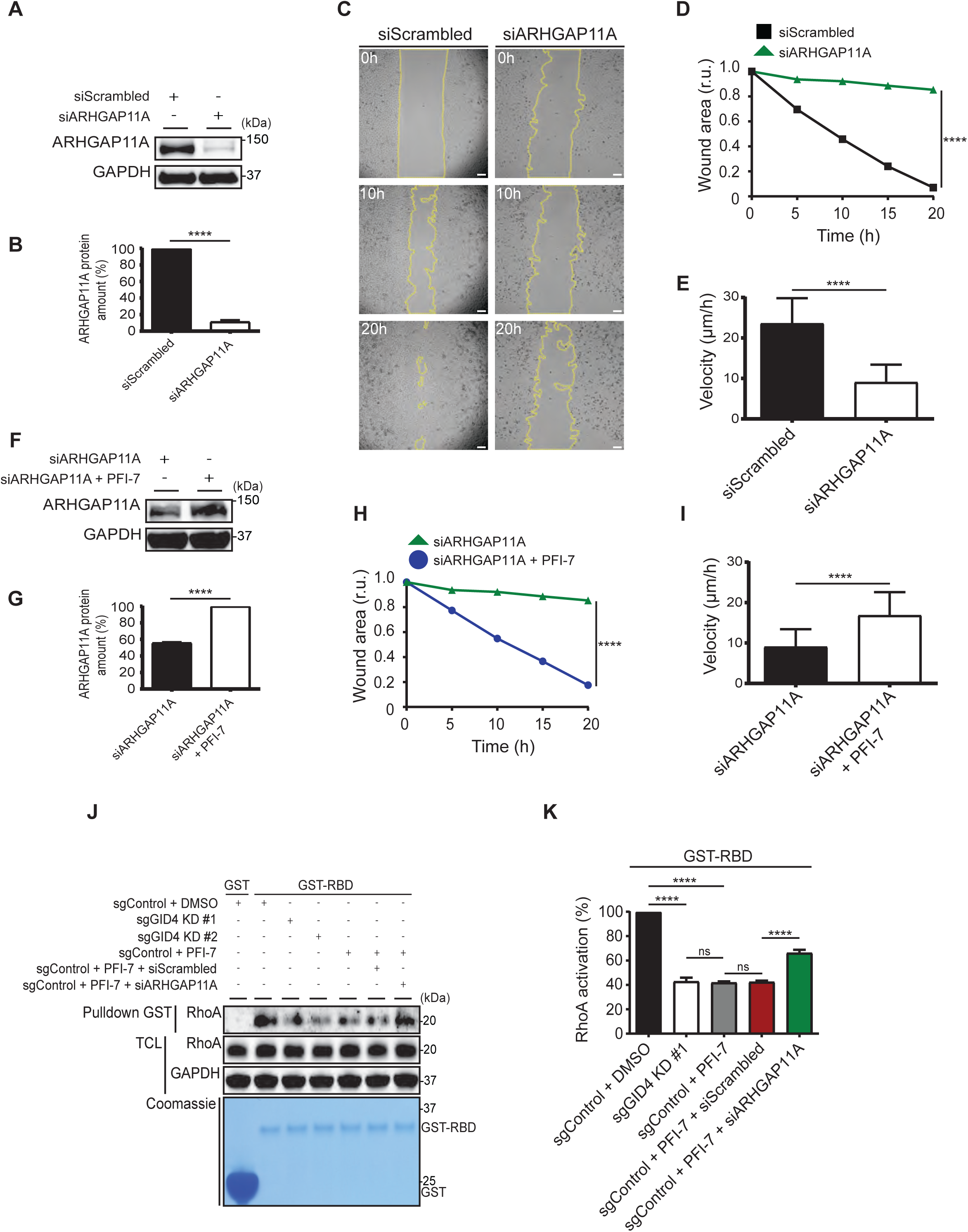
ARHGAP11A regulates cell migration downstream of GID4. A Western blots of sgControl HeLa total cell lysates (TCL) transfected with siScrambled (20 nM) or siARHGAP11A (20 nM). Blot of TCL was probed as indicated with antibodies to endogenous ARHGAP11A and GAPDH as a loading control. Data is representative of three independent experiments. B Bar graph from experiments as in Fig 4A showing percentage (%) of ARHGAP11A protein amount for sgControl HeLa cells transfected with siScrambled (20 nM) or siARHGAP11A (20 nM). Data values are normalized to siScrambled and are shown as mean ±SEM (n=3 independent experiments). The indicated *P*-values were calculated by a two-tailed Student’s *t* test. \*\*\*\**P*≤0.0001 C Representative brightfield images of sgControl HeLa cells transfected with siScrambled (20 nM) or siARHGAP11A (20 nM). Cells were grown to a monolayer with a defined cell-free gap establish by silicone insert. The silicone insert was removed, and images were acquired at 1 h intervals from initial time (0 h) to end (20 h). Images acquired at time 0 h, 10 h and 20 h are shown. The wound area was selected using the freehand selection tool (ImageJ) and is outlined in yellow. Scale bars, 100 µm. D Line graph for data generated as in Fig 4C showing wound area in relative units (r.u.) over time (h) for sgControl HeLa cells transfected with siScrambled (20 nM) or with siARHGAP11A (20 nM). Only data values at 20 hours were analyzed for statistical significance and are shown as mean ±SEM (n=3 independent experiments; 4 measurements were performed for each wounded area). The indicated *P*-values were calculated by a two-tailed Student’s *t* test. \*\*\*\**P*≤0.0001 E Bar graph of the velocity (µm/h) for data generated as in Fig EV3F with sgControl RPE1 cells transfected with siScrambled (20 nM) or with siARHGAP11A (20 nM). Data values are shown as mean ±SEM (n=3 independent experiments; 200 cells were analyzed for each condition). The indicated *P*-values were calculated by a two-tailed Student’s *t* test. \*\*\*\**P*≤0.0001 F Western blots of sgControl HeLa total cell lysates (TCL) transfected with siARHGAP11A (20 nM) or with siARHGAP11A (20 nM), and treated with PFI-7 (10 µM). Blot of TCL was probed with antibody as indicated to detect endogenous ARHGAP11A and GAPDH as a loading control. Images are representative of three independent experiments. G Bar graph from experiments as in Fig 4F showing percentage (%) of ARHGAP11A protein amount for sgControl HeLa cells transfected with siScrambled (20 nM) or with siARHGAP11A (10 nM), and treated with PFI-7 (20 µM). Data values are normalized to siScrambled and are shown as mean ±SEM (n=3 independent experiments). The indicated *P*-values were calculated by a two-tailed Student’s *t* test. \*\*\*\**P*≤0.0001 H Line graph showing the wound area in relative units (r.u.) over time (h) for data generated as in Figs 4C and EV3E, for sgControl Hela cells transfected with siARHGAP11A (20 nM) (the same data as in Fig 4D) or with siARHGAP11A (20 nM), and treated with PFI-7 (10 µM). Only data values at 20 hours were analyzed for statistical significance. Data values are shown as mean ±SEM (n=3 independent experiments; 4 measurements were performed for each wounded area). The indicated *P*-values were calculated by a two-tailed Student’s *t* test. \*\*\*\**P*≤0.0001 I Bar graph of migration velocity (µm/h) for data generated as in Fig EV3F of sgControl RPE1 cells transfected with siARHGAP11A (20 nM) or with siARHGAP11A (20 nM) and treated with PFI-7 (10 µM). Data values are shown as mean ±SEM (n=3 independent experiments; 200 cells were analyzed for each condition). The indicated *P*-values were calculated by a two-tailed Student’s *t* test. \*\*\*\**P*≤0.0001 J Pull-down assay in HeLa sgControl, sgGID4 KD #1, or sgGID4 KD #2 cells using GST tagged RBD domain of Rhotekin (GST-RBD) or GST alone (GST) as bait. Cells were treated as indicated with siScrambled (20 nM) or siARHGAP11A (20 nM) and were either untreated or treated with PFI-7 (10 µM). Upper panel, western blot with antibodies to RhoA was used to detect RhoA-GTP associated with the bait (Pulldown GST) or remaining in total cell lysate (TCL). Anti-GAPDH was used as a loading control. Lower panel, Coomassie gel shows GST or GST-RBD proteins. K Bar graph of RhoA-GTP protein amount (RhoA activation) as a percent (%) of total RhoA for the experiment as in Fig 4J. Data values are shown as mean ±SD (n=3 independent experiments). The indicated *P*-values were calculated by one-way ANOVA, followed by Bonferroni’s multiple comparisons test. ns (not significant, \*\*\*\**P*≤0.0001

Since ARHGAP11A functions as a RhoA-specific GAP to promote GTP hydrolysis, we examined whether altered RhoA activity may underlie the observed cell migration defects (Kagawa *et al*, 2013; Xu *et al*, 2013). To do this, we quantified RhoA-GTP levels by pull-down assay with GST fused-RBD domain of the RhoA-GTP effector Rhotekin (GST-RBD) (Figs 4J and K). Briefly, total cell lysates (TCL) prepared from GID4-depleted HeLa cell lines left untreated or treated with PFI-7 or DMSO were incubated with immobilized GST or GST-RBD to allow binding of active, RhoA-GTP. For control, we also analyzed RhoA-GTP levels in cells depleted for ARHGAP11A. The beads were washed, bound proteins eluted and bound RhoA-GTP immunoblotted with RhoA-specific antibodies. Interestingly, either GID4 depletion or PFI-7 treatment greatly reduced active RhoA compared to the DMSO-treated controls. However, ARHGAP11A knockdown by siRNA in PFI-7 treated sgControl cells significantly restored active RhoA levels when compared with PFI-7-treated sgControl cells (Figs 4J and K). Together, these results suggest that GID4 regulates cell motility by controlling active RhoA levels via ubiquitin-dependent degradation of ARHGAP11A. Since hyperactivation and inactivation of the RhoA-GTPase activity leads to similar migration-defects, we conclude that the concerted activity of ARHGAP11A and the hGID^GID4^ E3-ligase is required to maintain physiological levels and spatio-temporal dynamics of active RhoA during cell migration.

### ARHGAP11A accumulates at the cell periphery and colocalizes with F-actin protrusions in cells lacking GID4 activity

We used immunofluorescence to examine the subcellular localization of endogenous ARHGAP11A in HeLa cells exposed to different conditions. Consistent with previous results (Namba *et al*, 2020), ARHGAP11A localized to the cytoplasm and accumulated in nucleoli in sgControl cells (Fig 5A). siRNA-depletion or omission of the secondary antibody abolished this staining, demonstrating that the antibody specifically recognizes ARHGAP11A (Figs 5A and EV4). Interestingly, ARHGAP11A levels in the nucleus and cytoplasm increased upon PFI-7 treatment, with a particular accumulation at the cell periphery (Figs 5A and B), where it co-localized with F-actin-enriched protrusions (Figs 5A and B). A comparable phenotype was also observed in sgGID4-depleted cells (Fig 5C), with ARHGAP11A accumulating in nucleoli, the cytoplasm and to a lesser extent at the cell periphery (Figs 5C and D). In addition, GID4-depleted HeLa cells displayed a striking accumulation of actin stress fibers, and this phenotype was not altered by PFI-7 addition (Figs 5C and D). To quantify stress fiber anisotropy, we used the ImageJ FibrilTool (Boudaoud *et al*, 2014), attributing a score between 0 to define unordered, isotropic stress fibers and 1 for anisotropic stress fibers that are parallel and perfectly ordered. Indeed, sgGID4 depleted cells treated or not with PFI-7 showed an anisotropy score of 0.6, in contrast to 0.2 for sgControl cells (Fig 5D). The appearance of actin stress fibers was less pronounced in control cells treated with PFI-7 compared to GID4-depletion (Figs 5A-C), indicating that the GID4-inhibitor may only partially inhibit GID4 function under these conditions. Alternatively, GID4 may target additional substrates not dependent on the PFI-7-blocked binding pocket. We conclude that the hGID^GID4^ E3-ligase complex controls cell migration by degrading ARHGAP11A to prevent its accumulation at the cell periphery, thereby balancing RhoA activity.

**Fig 5.**
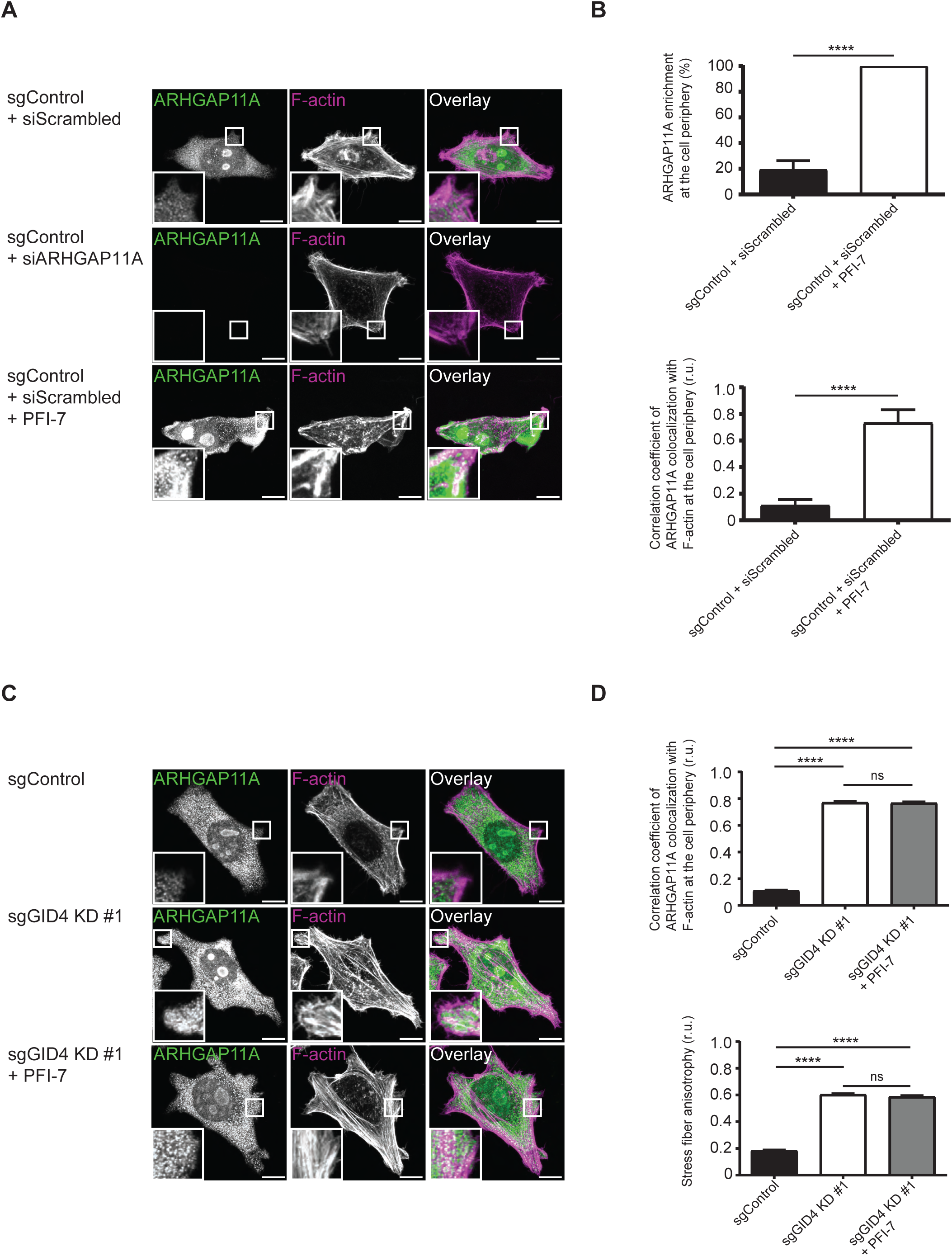
ARHGAP11A colocalizes with F-actin at cell periphery. A Representative confocal microscopy images showing immunofluorescence of ARHGAP11A with anti-ARHGAP11A antibody and staining of F-actin with phalloidin in sgControl HeLa cells. sgControl HeLa cells were transfected as indicated with siScrambled (20 nM) or siARHGAP11A (20 nM), and were untreated or treated with PFI-7 (10 µM). Green and magenta channels show overlayed merged images. The insets shown in lower left corner are magnified by factor 9. Images are representative of three independent experiments. Scale bars, 10 µm. B For cells transfected, and untreated or treated as in Fig 5A, the top bar graph shows enrichment in percentage (%) of endogenous ARHGAP11A within defined cell periphery regions, and the bottom bar graph shows the Pearson’s correlation coefficient data of the colocalization of endogenous ARHGAP11A with F-actin within defined cell periphery regions. Data values are shown as mean ±SEM (n=3 independent experiments; 100 cells were analyzed for each condition). The indicated *P*-values were calculated by a two-tailed Student’s *t* test. \*\*\*\**P*≤0.0001 C Representative confocal microscopy images showing immunofluorescence of ARHGAP11A with anti-ARHGAP11A antibody and staining of F-actin with phalloidin in sgControl or sgGID4 KD #1 HeLa cells with and without PFI-7 (10 µM) treatment. Green and magenta channels show overlayed merged images. The insets shown in lower left corner are magnified by factor 9. Images are representative of three independent experiments. Scale bars, 10 µm. D For cells treated as in Fig 5C, the top bar graph shows the Pearson’s correlation coefficient data of the colocalization of endogenous ARHGAP11A with F-actin within defined cell periphery regions, and the bottom bar graph shows stress fiber anisotropy calculated with ImageJ FibrilTool analysis (see materials and methods). Data values are shown as mean ±SD (n=3 independent experiments; 100 cells were analyzed for each condition). The indicated *P*-values were calculated by one-way ANOVA, followed by Bonferroni’s multiple comparisons test. ns (not significant), \*\*\*\**P*≤0.0001

## Discussion

Here we show that the hGID substrate receptor GID4 regulates cell growth and migration. Using a BioID2-approach, we identified several GID4 interactors and candidate substrates that use its conserved substrate-binding pocket, although only some of them comprise an N-terminal degron motif that fits the predicted criteria. Importantly, biochemical and functional characterization validated ARHGAP11A as a physiological hGID^GID4^ E3 ligase substrate. Indeed, ARHGAP11A is stabilized and accumulates at the cell periphery in cells with reduced GID4 levels and/or activity, leading to low RhoA-GTP levels and cell migration defects. Together, these results expand the cellular functions of hGID E3-ligase complexes and identify physiological substrates and mechanisms underlying specific phenotypes.

### The hGID^GID4^ E3 ligase regulates cell migration by degrading ARHGAP11A

Consistent with previous reports, our results suggest that the hGID E3 ligase complex triggers degradation of proteins involved in cell growth and migration (Maitland *et al*, 2022; Woo *et al*, 2012; Tripathi *et al*, 2015; Hasegawa *et al*, 2020; Ye *et al*, 2016). Indeed, loss of GID4 or inhibition of its binding pocket significantly impairs the wound healing response in HeLa and alters the motility of single RPE1 cells. Interestingly, we found that this defect is caused by GID4 interacting via its conserved substrate pocket with a cluster of proteins composed of ARHGAP11A, KIFC1, RACGAP1, and two other proteins associated with cytoskeleton organization or microtubule activity. ARHGAP11A functions as GTPase activating enzyme (GAP), targeting Rho-type GTPases involved in cytoskeletal organization, thereby regulating cell division, lymphocyte activation, myeloid leukocyte differentiation and leukocyte apoptosis. ARHGAP11A also regulates cell cycle progression by a RhoA-independent mechanism, as its depletion leads to cell cycle defects with high p27 levels (Lawson *et al*, 2016). Notably, while ARHGAP11A, KIFC1 and RACGAP1 are all turned-over with a half-life of less than 2 hours, only ARHGAP11A is stabilized in the absence of GID4 or upon PFI-7 treatment, indicating that KIFC1 and RACGAP1 are degraded *in vivo* by other pathways, possibly by the APC E3 ligase (Min *et al*, 2014).

Functionally, our results demonstrate that ARHGAP11A accumulation at the cell periphery of cells lacking GID4-activity leads to decreased levels of RhoA-GTP, resulting in migration defects. Conversely, ARHGAP11A depletion reduced cell proliferation and cell motility (Dai *et al*, 2018; Guan *et al*, 2021) by increasing RhoA activity. Indeed, GID4 inhibition by PFI-7 in ARHGAP11A-depleted cells partially restored ARHGAP11A protein levels and consequently cell motility, demonstrating that hGID^GID4^ antagonizes ARHGAP11A *in vivo*. Thus, both too much and too little RhoA activity interferes with cell motility and migration, consistent with the widespread spatio-temporal regulation of GTPases in organizing polarized actin structures and membrane protrusions. ARHGAP11A colocalizes with F-actin-enriched-protrusions and accumulates at the cell periphery of GID4-depleted cells, without apparent asymmetric localization and polarization. Surprisingly, GID4-depletion but not PFI-7 inhibition results in the formation of actin stress fibers, which are normally triggered by increased rather than decreased RhoA-GTP levels. Further work is now required to understand this apparent paradox. In addition to the cell periphery, ARHGAP11A is also enriched in nucleoli (Zanin *et al*, 2013; Namba *et al*, 2020), with unclear functional implications. Together, these results demonstrate that hGID^GID4^ regulates RhoA activity through ARHGAP11A and imply that ARHGAP11A steady-state levels need to be carefully balanced to allow directed cell migration. Although the regulatory mechanisms controlling GID4-mediated ARHGAP11A degradation under physiological conditions remain to be examined, we note that ARHGAP11A expression is increased in various cancers, including hepatocellular– and clear cell renal carcinoma and gastric cancer. Moreover, elevated ARHGAP11A levels have been associated with poor survival (Fan *et al*, 2021; Dai *et al*, 2018; Yang *et al*, 2023). It is thus tempting to speculate that loss of GID4 may increase ARHGAP11A stability in these cancer cells, thereby contributing to tumor progression.

### Beyond ARHGAP11A: Identification of additional GID4 substrates

Our comprehensive BioID2 screen identified over 20 GID4 substrate candidates that may be degraded by 26S proteasomes, highlighting the usefulness of the approach. As we included MG132 enrichment as a substrate criterion, we filtered out GID4 binding proteins whose ubiquitination may not lead to proteasomal degradation. For example, previous work revealed that GID4 binds several nucleolar RNA helicases including DDX17, DDX21, and DDX50 (Owens *et al*, 2023). While these targets rely on an N-terminal degron motif, their protein levels are not altered in the absence of GID4. Thus, the hGID^GID4^ complex may regulate cellular processes by degradative and non-degradative functions, and future work is needed to understand their selective ubiquitination mechanism.

In this study, GID4 substrates were defined by an increase in binding affinity upon MG132 treatment and dependence on a functional substrate-binding pocket. However, only a subset of them were also sensitive to PFI-7 treatment. The different interaction profiles of the drug and GID4^E237A^ mutant may originate from additional conformational changes in GID4^E237A^ besides disruption of its binding pocket. Consistent with this notion, we observed that the GID4-E237A mutant protein is less stable if cells are stressed with MG132, which may augment differences in the BioID2 analysis. Alternatively, PFI-7 competes with substrates binding, and thus affinity differences may contribute to the observed specificity profile.

Most of the identified GID4 substrate candidates are involved in chromatin organization, chromosome segregation and cell division, DNA binding and RNA-processing, and gene expression (Fig 2E), consistent with the nuclear localization of GID4. For example, we found that GID4 interacts with the Rb transcriptional corepressor 1 (RBL1), the cohesion subunit STAG2, the transcription factor ZNF106 and the ribosome-biogenesis regulator BYSL. While their GID4-dependent ubiquitination and proteasomal degradation remains to be determined, these candidates may contribute to the described function of GID4 in growth control and cell cycle regulation.

Surprisingly, only few putative GID4 substrates contain distinct N-terminal degron motifs that fulfill the stringent criteria proposed by earlier studies based on peptide assays. Besides ARHGAP11A, these targets include BYSL and ZNF106, which has been implicated in growth-related metabolism associated with early multiple organ failure (MOF) in acute pancreatitis (Grasberger *et al*, 2005; van den Berg *et al*, 2022). It is possible that some of the pocket-dependent GID4-interactors may bind indirectly, as exemplified by KIFC1, RACGAP1 and SHCBP1, known binding partners of ARHGAP11A (Fig 3A). Alternatively, the N-terminal degron may not be sufficient to mediate binding to GID4, or there might be GID4 substrates that utilize an interaction/degron motif that diverts from the proposed N-terminal structures. Finally, additional binding motifs may contribute to the interaction with GID4 and/or other subunits of the hGID complex. For example, we identified the E3-ligase subunit TRIP12 as a substrate candidate that binds GID4 in a pocket-dependent manner, but is also present in WDR26 pulldowns, suggesting that some substrates may interact with multiple GID subunits.

Finally, we identified a class of candidates that qualify as GID4 neo-substrates (Chana *et al*, 2022; Cowan & Ciulli, 2022), as these GID4 interactors accumulate in MG132-treated cells but exhibit increased binding in the presence of the PFI-7 inhibitor. These neo-substrates may bind GID4 through a binding site that gets preferentially exposed when the PFI-7 compound is blocking the conventional substrate pocket. Neo-substrates identification is an important factor to understand possible side effects of the PFI-7 compound, as exemplified by cereblon-induced degradation of the transcription factors Ikaros (IKZF1) and Aiolos (IKZF3), leading to severe embryonic defects in a drug-dependent manner (Krönke *et al*, 2014; Sievers *et al*, 2018).

### The BioID approach identified known and previously uncharacterized hGID subunits, along with potential regulators

The BioID2 screen also allowed to identify interactors that are unlikely to be GID4 substrates but rather function as hGID subunits or regulators (Fig 2B). Indeed, the interaction of these GID4 binding proteins was not altered by MG132-treatment, and they similarly bound both wild-type and the GID4-E237A substrate-pocket mutant. Likewise, the interaction was not diminished by PFI-7 treatment. Indeed, our GID4 BioID2 screen identified multiple known hGID subunits, including RANBP9 and RANBP10, which bind the hGID complex by a mutually exclusive mechanism. In contrast, we did not find a human homologue of GID12, a GID4-interacting protein that sterically blocks substrate ubiquitination (Qiao *et al*, 2022). However, we reveal previously unidentified hGID candidate subunits and/or regulators, including CHD9, GPATCH4, WIZ or INTS12. GPATCH4 is predicted to regulate cell growth and differentiation of hematopoietic progenitor cells, a function reminiscent of the requirements of the hGID complex in erythroid maturation (Hirawake-Mogi *et al*, 2021). CHD9, WIZ and INTS12 are nuclear proteins involved in gene expression or chromatin regulation (Alendar *et al*, 2020; Simon *et al*, 2015; Isbel *et al*, 2016; Baillat *et al*, 2005) and may thus help recruiting the hGID complex to regulate DNA-associated processes and/or polymerase activity. Further work is required to validate these candidates, but their functional relevance is coherent with the identified GID4 substrates. It will be interesting to extend the BioID approach to other hGID core subunits, for example to find mammalian GID4-like substrate receptors that, analogous to yeast, are recruited into the hGID complex via ARMC8 (Langlois *et al*, 2022; Kong *et al*, 2021). Moreover, BioID screens using the alternative hGID substrate receptors WDR26 and MKLN1 as baits may reveal new classes of hGID substrates and degron motifs.

### BioID: a valid approach to identify E3-ligase substrates and functions

The identification of physiological E3 ligase substrates is often hampered by the generally low affinity of substrate-receptor interactions that cannot easily withstand cell lysis and stringent immunoprecipitation conditions. To circumvent this bottleneck, BioID screening emerged as an alternative approach, as biotinylation of interacting proteins is dictated by their close proximity in cells prior to lysis and extract preparation (Roux *et al*, 2012). Until recently, BioID approaches suffered from severe specificity limitations, fueled by the need to overexpress bulky fusion proteins combined with long incubation times to reach sufficient labeling. However, the recent development of smaller biotinylation enzyme variants with increased catalytic activity (for example miniTurboID, UltraID) mitigated some of these risks (Branon *et al*, 2018; Kubitz *et al*, 2022). Nevertheless, including secondary filtering criteria such as the presence of motifs and/or domains, or stringent specificity controls such as treatment with MG132 substantially reduce unspecific noise of BioID measurements. Ideally, inhibitory compounds or mutant proteins altering binding to critical components, for example mutations in the substrate interaction domain, further help to distinguish direct from indirect, unspecific interactors. Additionally, rapid improvements in Alphafold to predict binding surfaces with atomic accuracy using a deep learning algorithm greatly facilitates the identification of critical residues (Jumper *et al*, 2021). As shown by this and other studies (Sharifi Tabar *et al*, 2022), including such specificity criteria allows for efficient filtering of comprehensive BioID data sets, making this approach a valid alternative to more conventional methods such as AP-MS and diGly enrichments (Iconomou & Saunders, 2016). Therefore, advanced BioID screening strategies hold great potential to study other multi-subunit RING-domain-containing E3 ligases, particularly those for which physiological substrates remain scarce despite known substrate receptors and/or functions.

### Materials and Methods Reagents and Tools

Experimental models, plasmids, antibodies, oligonucleotides, chemicals and other reagents, and softwares used in this study are listed in the Reagents and Tools Table.

### Cell culture, siRNA transfections, and generation of stable Flp-In T-REx cell lines

Cells were cultured in DMEM supplemented with 10% fetal bovine serum and 1% penicillin-streptomycin (DMEM\FBS\PS) and maintained at 37°C in 5% CO_2_. For siRNA transfections, a final concentration of 20 nM ON-TARGETplus Non-targeting Pool (siScrambled) or 20 nM ON-TARGETplus Human ARHGAP11A siRNA was transfected in the presence of Lipofectamine 2000 according to the manufacturer’s recommendations. Stable Flp-In T-REx HeLa cell lines expressing BirA2-Flag-GID4, BirA2-Flag-GID4^E237A^, BirA2-Flag-EGFP or an empty vector were generated as described elsewhere (Kean *et al*, 2012; Bagci *et al*, 2020). Protein expression and biotinylation were induced in the presence of 1 µg/ml tetracycline and 50 µM biotin.

### Generation of CRISPR/BAC GID4 KD HeLa or RPE1 cell lines

The clustered regularly interspaced short palindromic repeat (CRISPR)-Bac cells were generated as described elsewhere, with modifications (Schertzer *et al*, 2019). We designed four single guide RNAs (sgRNAs) targeting different exons of the *Gid4* gene. Each sgRNA was separately cloned into the PB_rtTA_BsmBI vector to generate the following vectors: PB_rtTA_Bsmb1_Gid4_sgRNA1, PB_rtTA_Bsmb1_Gid4_sgRNA2, PB_rtTA_Bsmb1_Gid4_sgRNA3 and PB_rtTA_Bsmb1_Gid4_sgRNA4. As RPE1 cells are resistant to Hygromycin B, the Hygromycin B-resistance (HygR) gene in the PB_tre_Cas9 vector was replaced with a Puromycin-resistance gene (PuroR) using the NEBuilder HiFi DNA assembly kit to generate the PB_tre_Cas9_puro vector. The HiFi reaction was performed according to the manufacturer’s recommendations. We then simultaneously cotransfected 625 ng of either PB_tre_Cas9 (containing HygR) or PB_tre_Cas9_puro (containing PuroR) with 1250 ng of the Super piggyBac Transposase expression vector and 157 ng of each of the four PB_rtTA_BsmBI vectors with sgRNAs targeting the *Gid4* gene. As a negative control, we cotransfected the empty pb_rtTA_Bsmb1 vector into which we did not clone a sgRNA-targeting sequence (control). HeLa cells were selected in the presence of Hygromycin B (200 µg/ml) and G418 (200 µg/ml) for 10 days. RPE1 cells were selected in the presence of Puromycin (10 µg/ml) and G418 (200 µg/ml) for 10 to 20 days. Cell death was observed within 3 or 4 days upon G418 and Hygromycin B or Puromycin treatment. After selection, cells were cultured in the absence of G418 and Hygromycin B or Puromycin, and doxycycline (1 µg/ml) was added to the DMEM\FBS\PS medium for 4 days to induce the Cas9 expression. Efficiency of GID4 knockdown was assessed by western blot.

### MTT assay

3-(4,5-dimethylthiazol-2-yl)-2,5-diphenyl-2H-tetrazolium bromide (MTT) assays were carried out as described elsewhere, with modifications (Stier *et al*, 2023). The MTT assay kit (Promega) was used to measure the cell proliferation according to the manufacturer’s recommendations. 5000 HeLa cells were plated in 96 well plates with three biological replicates per condition. Cells were grown in 96 well plates for 24 h, 48 h or 72 h prior to the incubation with the MTT dye master mix for 2 h at 37°C in 5% CO_2_. The reaction was stopped by adding 100 µl stop solution. Plates containing the MTT-treated cells were measured at a wavelength of 570 nm and the growth rate was normalized to day 0.

### Wound healing

Two-well silicone inserts with a defined cell-free gap (Ibidi) were inserted into 8 well microscopy slides (Ibidi). For each condition, 10000 HeLa cells were plated into each chamber and incubated overnight at 37°C in 5% CO_2_ in the presence or absence of DMSO (10 µM) or PFI-7 (10 µM). For siRNA transfections, siScrambled-or siARHGAP11A-transfected cells were plated into chambers after 24 h of siRNA transfections. Next day, medium was replaced with or without DMSO (10 µM) or PFI-7 (10 µM) and placed inside the CO_2_ incubator of a phase-contrast microscope. Time-lapse imaging was performed by taking an image every 1h for 24h using 10X objective. 3 random fields were acquired for each of the three biological replicates. The Nikon Ti2-E widefield microscope equipped with a xy stage (Prior), a piezo z-drive (Prior) and the NIS-Elements software were used to take images. Captured images were analyzed using the ImageJ software. The size of the gap area at times 0, 5, 10, 15 or 20 h was measured using the freehand selection tool and analyzed using the Measure command in the Analyze menu. The measured gap area at times 5, 10, 15 or 20 h was normalized to 0 h, to determine the wound area closure in relative units.

### Single-cell tracking

Single-cell tracking was performed and analyzed as described elsewhere, with modifications (Pijuan *et al*, 2019). 5000 RPE1 cells per condition were plated into 8 well microscopy slides (Ibidi) and incubated overnight at 37°C in 5% CO_2_ in the presence or absence of DMSO (10 µM) or PFI-7 (10 µM). Like wound healing assays, siScrambled-or siARHGAP11A-transfected cells were plated into chambers after 24 h of siRNA transfections. Next day, medium was replaced with or without DMSO (10 µM) or PFI-7 (10 µM) and placed inside the CO_2_ incubator of a phase-contrast microscope. Time-lapse imaging was performed by taking an image every 30 min for 24 h using 10X objective. 8 random fields were acquired for each of the three biological replicates to obtain data from at least 200 cells. The Nikon Ti2-E widefield microscope equipped with a xy stage (Prior), a piezo z-drive (Prior) and the NIS-Elements software were used to take images. Captured images were analyzed using the manual tracking plugin and chemotaxis tool (Ibidi) in the ImageJ software. The tracking plot, velocity and overlay dots and lines data were generated as described elsewhere (Pijuan *et al*, 2019).

### BioID2 and MS

BioID2-MS experiments were carried out as previously described, with modifications (Bagci *et al*, 2020; Methot *et al*, 2018; Couzens *et al*, 2013; Mehnert *et al*, 2020). BirA2-Flag-expressed cells were harvested after 24 h of tetracycline, biotin, MG132, DMSO or PFI-7 treatment. They were washed three times in Phosphate-buffered saline (PBS) and lysed in 1.5 ml radio-immunoprecipitation assay (RIPA) buffer. They were then sonicated 30 s at 30% amplitude (three times 10 s bursts with 2 s break between). 1 µl benzonase added to each sample followed by a centrifugation for 30 min at 4 °C at maximum speed. Cleared lysates were incubated with 70 µl of streptavidin beads at 4°C for 3 h with rotation. Streptavidin beads were then transferred to chromatography columns, washed three times with lysis buffer, then three times with 50 mM ammonium bicarbonate (ABC), pH 8.0. Samples were resuspended in 200 µL ABC, transferred to centrifugal units, centrifuged for 15 min at 4°C at 8000 G. 100 µl of 8M urea and 1µl of 500 mM tris(2-carboxyethyl)phosphine (TCEP) were added to each sample, incubated for 30 min at 37°C at 600 rpm. 2 µl of 500 mM iodoacetamide was added to each sample, incubated for an additional 30 min at 37°C at 600 rpm. Samples were then centrifuged for 15 min at 8000 G and washed two times in 200 µl ABC. 100 µl ABC and 2.5 µl trypsin were added to each sample prior to incubation for 12 h at 37°C at 700 rpm. Next day, Tryptic proteolysis was quenched with 5% formic acid and peptides were subjected to C18 cleanup, according to the manufacture’s recommendations. Eluted peptides were dried using a speed vacuum and resuspended in 20ul of 2% acetonitrile and 0.1% formic acid. LC-MS/MS analysis was performed on an Orbitrap QExactive + mass spectrometer (Thermo Fisher) coupled to an EASY-nLC-1000 liquid chromatography system (Thermo Fisher). Peptides were separated using a reverse phase column (75□µm ID x 400□mm New Objective, in-house packed with ReproSil Gold 120 C18, 1.9 µm, Dr. Maisch GmbH) across 120□min linear gradient from 5 to 40% (buffer A: 0.1% (v/v) formic acid; buffer B: 0.1% (v/v) formic acid, 95% (v/v) acetonitrile). The DDA data acquisition mode was set to perform one MS1 scan followed by a maximum of 16 scans for the top 16 most intense peptides (TOP16) with MS1 scans (R=70’000 at 400□m/z, AGC=1e6 and maximum IT=100 ms), HCD fragmentation (NCE=27%), isolation windows (2.0 m/z) and MS2 scans (R=17’500 at 400□m/z, AGC=1e5 and maximum IT=50ms). A dynamic exclusion of 30s was applied and charge states lower than two and higher than seven were rejected for the isolation.

### MS data analyses

MS data analyses were performed as described previously (Bagci *et al*, 2020). RAW files were converted to .mzXML files using the Proteowizard tool version 3.0.19014 (Kessner *et al*, 2008). Peptide search and identification were carried out by using Human RefSeq Version 57, mascot search engine and the iProphet pipeline included in ProHits (Liu *et al*, 2012; Shteynberg *et al*, 2011). Mascot parameters were determined with trypsin specificity allowing up to 2 missed cleavages. Carbamidomethyl (C) was used as a fixed modification. Oxidation (M) and deamidation (NQ) were enabled as variable modifications. Mass tolerances for precursor and fragment ions were used as 15 ppm and 0.6 Da, respectively. Peptide charges were set to +2, +3 and +4. Mascot search results were processed using PeptideProphet and peptide assembly into protein was performed using Trans-Proteomic Pipeline (Deutsch *et al*, 2023).

### Dot plot analyses

SAINT output files of untreated or MG132 and/or PFI-7-treated BirA2-Flag-GID4, BirA2-Flag-GID4^E237A^ bait data analyzed in ProHits were processed using the ProHits-viz platform to carry out dot plot analyses (Knight *et al*, 2017; Choi *et al*, 2011). Experimental controls such as the empty pcDNA5-vector and BirA2-Flag-EGFP treated or not with MG132 have been used in SAINT analyses to filter non-specific interactions.

### GO term analyses

Gene Ontology (GO) term analyses were performed as described previously, with modifications (Shooshtarizadeh *et al*, 2019). The g:Profiler tool was used to analyze the functionally annotated GO terms (Reimand *et al*, 2016). Molecular function or biological process of the prey proteins identified in the MG132-treated BirA2-Flag-GID4, but not in untreated BirA2-Flag-GID4, are shown in heat map analyses. Preys identified in SAINT output files were submitted to g:Profiler for GO term analysis. The GO term enrichment score is displayed as the –log10 of adjusted *P* values.

### Protein-protein association networks and clustering

Proteins identified in SAINT output files were submitted to the STRING database to generate the protein-protein association networks or functionally relevant protein clusters. Protein clusters were obtained following MCL clustering using STRING v11.5 (Szklarczyk *et al*, 2019). The MCL inflation parameter is 3 and the protein-protein interaction (PPI) enrichment p-value is 2.33×10^-12^. Known interactions were extracted from curated databases including Biocarta, BioCyc, GO, KEGG or Reactome. Experimentally determined interactions were extracted from BIND, DIP, GRID, HPRD, IntAct, MINT or PID. Text mining-based interactions were extracted from the scientific literature as determined by STRING (Szklarczyk et al, 2019).

### Protein sequence alignment and analysis

Alignment and analysis of prey protein sequences identified in BioID2 were performed using the Jalview software vs 2.11.2.6 (Waterhouse *et al*, 2009). The first N-terminal amino acids, excluding Met at the first position, were submitted to Jalview. The conserved residues were colored by using the clustal color scheme.

### Co-immunoprecipitations, GST-RBD pulldown, ubiquitination assays and half-life measurements

#### Co-immunoprecipitations and Ubiquitination assay

Flp-In T-REx HeLa cells from experiments as in (Figs 2H and EV2B) were lysed in 3-((3-cholamidopropyl) dimethylammonio)-1-propanesulfonate (CHAPS) buffer (30 mM Tris-HCl, pH 7.5, 150 mM NaCl, 5 mM MgCl_2_ and 1% CHAPS) supplemented with inhibitors (5 mM NaF, 1 mM Na_2_VO_4_ and 1x complete, EDTA-free Protease Inhibitor Cocktail). HeLa cells from experiments as in (Fig 3F) were lysed in Nonidet P40 (NP40) buffer (30 mM Tris-HCl, pH 7.5, 150 mM NaCl, 5 mM MgCl_2_ and 1% NP40) supplemented with the same inhibitors used in experiments as in (Fig 2H). In both experiments, cleared total cell lysates (TCL) were incubated for 3 h at 4°C with anti-flag M2 affinity gel (FLAG beads). Bound co-immunoprecipitated proteins (IP-FLAG) and unbound total cell lysates (TCL) were subjected to western blotting using the indicated antibodies against endogenous or exogenous (FLAG, Myc or HA-tagged) proteins.

#### GST-RBD pulldown

GST or GST-RBD fusion proteins were purified from BL21 bacteria and coupled with GST beads as described previously (Bagci *et al*, 2020). Expression of purified GST or GST-RBD were assessed by Coomassie. HeLa sgControl, sgGID4 KD #1 or sgGID4 KD #2 cells from experiment as in (Fig 4J) were transfected with siScrambled (20 nM) or siARHGAP11A (20 nM) and were either untreated or treated with PFI-7 (10 µM). Cells were then lysed in CHAPS buffer with the aforementioned inhibitors and incubated with GST beads coupled with GST or GST-RBD for 3 h at 4°C. Bound pulldown proteins (pulldown GST) and unbound total cell lysates (TCL) were subjected to western blotting using the indicated antibodies against endogenous RhoA or GAPDH proteins.

#### Half-life measurements

HeLa sgControl or sgGID4 KD #1 cells from experiments as in (Figs 3B-E) were untreated (time at 0 h) or treated with 20 µg/ml cycloheximide (CHX) at times 2 h, 4 h or 8 h. Cells were lysed in CHAPS buffer with the aforementioned inhibitors. Extracted total cell lysates (TCL) were subjected to western blotting using the indicated antibodies against endogenous ARHGAP11A, KIFC1, RACGAP1 or GAPDH.

### Immunofluorescence

7500 HeLa cells per condition were plated on coverslips one day prior to fixation with 4% paraformaldehyde (PFA). Cells were permeabilized with PBS/0.30% Triton-X100, blocked in PBS/0.30% Triotin-X100/1% bovine serum albumin (BSA) for 1 h at RT followed by an overnight incubation with the ARHGAP11A antibody, or not. Next day, samples were washed three times in PBS and incubated with secondary anti-rabbit IgG Alexa Fluor 488 antibody for 1 h at RT with 1:1000 dilution. Samples were then washed three times in PBS and incubated with rhodamine phalloidin for 1 h at RT with 1:400 dilution. Samples were washed again three times with PBS, incubated with DAPI for 5 min at RT with 1:10000 dilution. Samples were washed three times with PBS. Cover slips were then mounted on microscopy slides using ProLong Diamond Antifade Mountant and fixed with a nail polish. The Leica SP8-AOBS confocal microscope was used to take images, using a 63x/1.4 oil immersion objective. The following excitation lasers were used: 405 nM for DAPI (blue channel), 488 nM for Alexa Fluor 488 (green channel) and 561 nM for rhodamine phalloidin (magenta channel). Images were processed using the Leica Application Suite X (Las X) software. Contrast was adjusted throughout the whole image to enhance visibility when necessary. Z-stack images were converted to maximum projections and exported to Adobe Illustrator to prepare figures. 100 cells were analyzed for each condition. To quantify percentage of ARHGAP11A enrichment at the cell periphery, regions of interest (ROIs) were created after finding the boundaries of cell periphery by setting a threshold on the green channel, as described elsewhere (DesMarais *et al*, 2019). Nuclear fluorescence of ARHGAP11A was excluded from the analysis. Fluorescence intensity of the ROIs was analyzed using the ImageJ tool and exported to the GraphPad Prism 9 software for further analysis. The BIOP JACoP plugin integrated in ImageJ was used to quantify the colocalization of ARHGAP11A with F-actin at the cell periphery. Region of interests (ROIs) surrounding the cell periphery were selected on green and magenta channels. Auto-thresholding using the Otsu method was applied for each colocalization. The output data showing the Pearson correlation coefficient were transferred to the GraphPad Prism 9 software for analysis. Stress fiber anisotropy was quantified using the FibrilTool plugin in ImageJ as described elsewhere (Boudaoud et al., 2014). The corresponding channel for each image was selected for the fiber array signals and the value for “Multiply line length by” set to the default value 1. The Polygon tool was used to select the ROIs by eye displaying stress fibers prior to the FibrilTool analysis. A score between 0 (unordered, isotropic stress fibers) and 1 (perfectly ordered, parallel, anisotropic stress fibers) was used to analyze the stress fiber anisotropy.

### Data presentation and statistical analysis

Bayesian False Discovery Rate (BFDR) of 0.01 was used to filter non-specific BioID2-MS interactions from the dot plot analyses. The GO term enrichment score was determined as the –log10 of adjusted *P* values, calculated by the g:Profiler tool. For other experiments, GraphPad Prism 9 was used to generate quantification graphs and carry out statistical analysis. Co-immunoprecipitation, GST-RBD pulldown or half-life measurements were quantified using the ImageJ software. Selected lanes (Analyze/Gels/Plot Lanes) were plotted, and the intensity of each protein band was measured using the wand tracing tool and normalized to the total RhoA (for GST-RBD pulldown) or GAPDH levels (for all other western blot experiments). For experiments including multiple conditions, the indicated *P*-values were calculated using one-way ANOVA, followed by Bonferroni’s multiple comparisons. Data values for experiments comparing multiple conditions are shown as mean ±standard deviation (SD). For experiments comparing two conditions, the indicated *P*-values were calculated using a two-tailed Student’s *t* test. Data values for experiments comparing two conditions are shown as mean ±standard error of the mean (SEM). The indicated P values are as follows: ns (not significant), \**P*≤0.05, \*\**P*≤0.01, \*\*\**P*≤0.001, \*\*\*\**P*≤0.0001. Figs 1A and 2A were created using the BioRender software.

## Author Contributions

**Halil Bagci**: Conceptualization; data curation; formal analysis; investigation; methodology; project administration; software; validation; visualization; writing-original draft. **Martin Winkler**: Data curation; formal analysis; investigation; methodology. **Federico Uliana**: Data curation; formal analysis; investigation methodology; software. **Jonathan Boulais**: Data curation; formal analysis. **Weaam I Mohamed**: Data curation; formal analysis; investigation; methodology; **Sophia L Park**: Data curation; formal analysis; investigation; methodology; **Jean-François Côté**: Funding acquisition; resources; Supervision. **Matthias Peter**: Conceptualization; funding acquisition; methodology; project administration; resources; supervision; visualization; writing-original draft. All authors contributed to the editing of the manuscript.

## Acknowledgments

We thank the ScopeM facility members, Tobias Schwarz and Joachim Hehl for microscopy training and technical assistance and Gabor Csucs for helpful discussion with single-cell tracking experiments. We are grateful to Anne-Claude Gingras for the BioID2 plasmids, Brenda A. Schulman for the GID4 antibody, and Stephen Taylor for the Flp-In T-REx HeLa cell line. PB_rtTA_BsmBI and PB_tre_Cas9 plasmids were provided by Mauro Calabrese (Addgene plasmids # 126028 and 126029), GST-RBD by Martin Schwartz (Addgene plasmid # 15247) and pCMV6-AN-HA_Ubiquitin was a gift from Roger Woodgate (Addgene plasmid # 131258). We thank Gabriel Neurohr, Arun Peter and Frank vanDrogen for critical feedback, Alicia Smith for editing of the manuscript, Christian Poitras for MS data management and members of the Peter lab for helpful discussions. This work was supported by the Swiss National Science Foundation (310030_179283/1) to Matthias Peter, and the National Science and Engineering Research Council of Canada (RGPIN-201604808) to Jean-François Côté.

## Figure legends

**Fig EV1.** Abrogation of GID4 causes cell migration defect. A Western blots of RPE1 total cell lysates (TCL) indicating GID4 and WDR26 protein expression. Lysates are derived from a stable clone without sgRNA (sgControl), a stable clone with a pool of four sgRNAs targeting *Gid4* (sgGID4 KD #1), a second stable clone with a pool of four sgRNAs targeting *Gid4* (sgGID4 KD #2), sgControl treated with DMSO (10 µM), sgControl treated with PFI-7 (10 µM), sgGID4 KD #1 treated with PFI-7 (10 µM) or sgGID4 KD #2 treated with PFI-7 (10 µM). Blot shown is representative of three independent experiments. B Additional quantification of the MTT assay shown in Fig 1C. HeLa cells measuring absorbance at 570nm to indicate cell metabolic activity during 1, 2 or 3 days (d) for lysates derived as in (B). HeLa sgControl, sgGID4 KD #1, or sgGID4 KD #2 cells were untreated or treated with DMSO (10 µM) and/or PFI-7 (10 µM). Data values at day 3 were analyzed for statistical significance and are shown as mean ±SD (n=3 independent experiments; 3 biological replicates were performed for each experiment). The indicated *P*-values were calculated by one-way ANOVA, followed by Bonferroni’s multiple comparisons test. ns (not significant), \**P*≤0.05, \*\*\*\**P*≤0.0001 C Representative brightfield images acquired over time (h) of a wound healing assay with HeLa sgGID4 KD #2 cells, either untreated or treated with PFI-7 (10 µM). Cells were grown to a monolayer with a defined cell-free gap established by silicone insert. The silicone insert was removed, and images were acquired at 1 h intervals from initial time (0 h) to end (20 h). Images acquired at time 0 h, 10 h and 20 h were shown. Wound area was selected using the freehand selection tool (ImageJ) and is outlined in yellow. Scale bars, 100 µm D Bar graph showing wound area measurements in relative units (r.u.) from images acquired as in Figs 1D and EV1C with wound area selected as in Fig 1D and EV1C and normalized to starting cell-free gap area at time 0h. Data values at 20 hours were analyzed for statistical significance and are shown as mean ±SD (n=3 independent experiments; 4 measurements were performed for each wounded area). The indicated *P*-values were calculated by one-way ANOVA, followed by Bonferroni’s multiple comparisons test. ns (not significant), \*\*\*\**P*≤0.0001 E Representative brightfield images showing a positing of individual RPE1 cells at the initial time (0h) and final time (24 h), with a cell trajectory (colored lines, 24 h) generated from merged individual cell localization over the 24 h period. Images were acquired at 30 min intervals for 24h. RPE1 sgControl, sgGID4 KD #1 or sgGID4 KD #2 cells were either untreated or treated with DMSO (10 µM) or PFI-7 (10 µM), as indicated. Scale bars, 100 µm. F Plots showing a 24 h period of merged individual RPE1 cell trajectories set to a common origin at the intersection of y (µm) and x (µm) axes for sgControl and sgGID4 KD #2 cells, either untreated or treated with DMSO (10 µM) or PFI-7 (10 µM). Images were acquired at 30 min intervals for 24 h, analyzed using a manual tracking plugin and chemotaxis tool (Ibidi) in ImageJ software. G Bar graph showing cell velocity (µm/h) of RPE1 cells from data acquired and analyzed as in Figs EV1E and F. Data values are shown as mean ±SD (n=3 independent experiments; 200 cells were analyzed for each condition). The indicated *P*-values were calculated by one-way ANOVA, followed by Bonferroni’s multiple comparisons test. ns (not significant), \*\*\*\**P*≤0.0001

**Fig EV2.** BioID2-GID4 constructs are functional. A Western blots of total cell lysate (TCL) generated from Flp-In T-REx HeLa cell lines expressing either BirA2*-Flag-EGFP, BirA2*-Flag-GID4, or BirA2*-Flag-GID4E^237A^, either untreated or treated with PFI-7 (10 µM). Expression and biotinylation of endogenous proteins were determined using anti-Flag-HRP and anti-Streptavidin-HRP antibodies, respectively. Anti-GAPDH was used as a loading control. BirA2*-Flag was used as the empty vector control. B Western blots of bound co-immunoprecipitated proteins (IP-FLAG) and unbound total cell lysate (TCL) from Flp-In T-Rex HeLa cell lines expressing BirA2*-FLAG, BirA2*-FLAG-GID4, or BirA2*-FLAG-GID4^E237A^ as bait proteins. Blots were probed where indicated with antibodies to FLAG or to endogenous MAEA, and GAPDH as a loading control.

**Fig EV3.** ARHGAP11A mediates migration downstream of GID4. A Western blots of sgControl RPE1 total cell lysates (TCL) transfected with siScrambled (20 nM) or siARHGAP11A (20 nM). Blot of TCL was probed as indicated with anti-ARHGAP11A antibody to detect endogenous ARHGAP11A protein and anti-GAPDH antibody as a loading control. Data is representative of three independent experiments. B Bar graph from experiments as in Fig EV3A showing percentage (%) of ARHGAP11A protein amount for sgControl RPE1 cells transfected with siScrambled (20 nM) or with siARHGAP11A (20 nM). Data values are normalized to siScrambled and are shown as mean ±SEM (n=3 independent experiments). The indicated *P*-values were calculated by a two-tailed Student’s *t* test. \*\*\*\**P*≤0.0001 C Western blots of sgControl RPE1 cell lysates (TCL) transfected with siARHGAP11A (20 nM) or with siARHGAP11A (20 nM) in the presence of PFI-7 (10 µM). Blot of TCL was probed as indicated with anti-ARHGAP11A antibody to detect endogenous ARHGAP11A protein and anti-GAPDH antibody as a loading control. Images are representative of three independent experiments. D Bar graph from experiments as in Fig EV3C showing percentage (%) of ARHGAP11A protein amount for sgControl RPE1 cells transfected with siScrambled (20 nM) or with siARHGAP11A (20 nM) in the presence of PFI-7 (10 µM). Data values are normalized to siScrambled and are shown as mean ±SEM (n=3 independent experiments). The indicated *P*-values were calculated by a two-tailed Student’s *t* test. \*\*\*\**P*≤0.0001 E Representative brightfield images acquired over time (h) of a wound healing assay with sgControl HeLa cells transfected with siARHGAP11A (20 nM) and PFI-7 (10 µM). Cells were grown to a monolayer with a defined cell-free gap established by silicone insert. The silicone insert was removed, and images were acquired at 1 h intervals from initial time (0 h) to end (20 h). Images acquired at time 0 h, 10 h and 20 h were shown. Wound area was selected using the freehand selection tool (ImageJ) and is outlined in yellow. Scale bars, 100 µm F Plots showing a 24 h period of merged individual sgControl RPE1 cell trajectories set to a common origin at the intersection of y (µm) and x (µm) axes. Cells were transfected with siScrambled (20nm) or siARHGAP11A (20 nM), with or without PFI-7 (10 µM). Images were acquired at 30 min intervals for 24 h and analyzed using a manual tracking plugin and chemotaxis tool (Ibidi) in ImageJ software. G Representative brightfield images showing positions of individual sgControl RPE1 cells at the initial time (0h) and final time (24 h), with a cell trajectory (colored lines, 24 h) generated from merged individual cell localization over the 24 h time period. Cells were transfected with siScrambled (20nm) or siARHGAP11A (20 nM), with or without PFI-7 (10 µM). Images were acquired at 30 min intervals for 24h. Scale bars, 100 µm.

**Fig EV4.** Secondary antibody staining for immunofluorescence. A Representative confocal microscopy images showing immunofluorescence when only secondary anti-ARHGAP11A antibody is used, with staining of F-actin with phalloidin in sgControl HeLa cells. Green and magenta channels show overlayed merged images. The insets shown in lower left corner are magnified by factor 9. Images are representative of three independent experiments. Scale bars, 10 µm.

