## Supplementary figures and images for "The hGID^GID4^ E3 ubiquitin ligase complex targets ARHGAP11A to regulate cell migration"

### Figure EV1

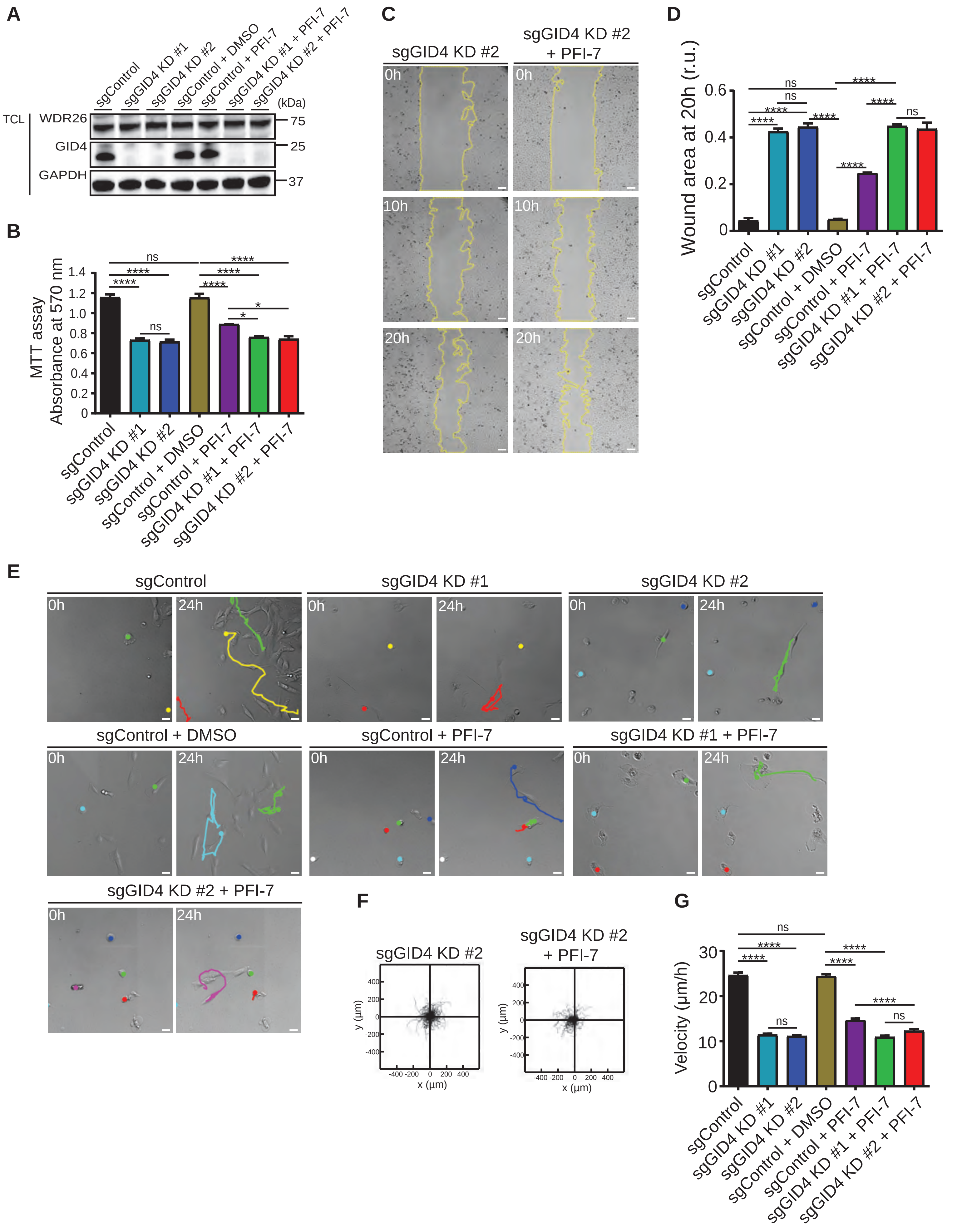

### Figure EV2

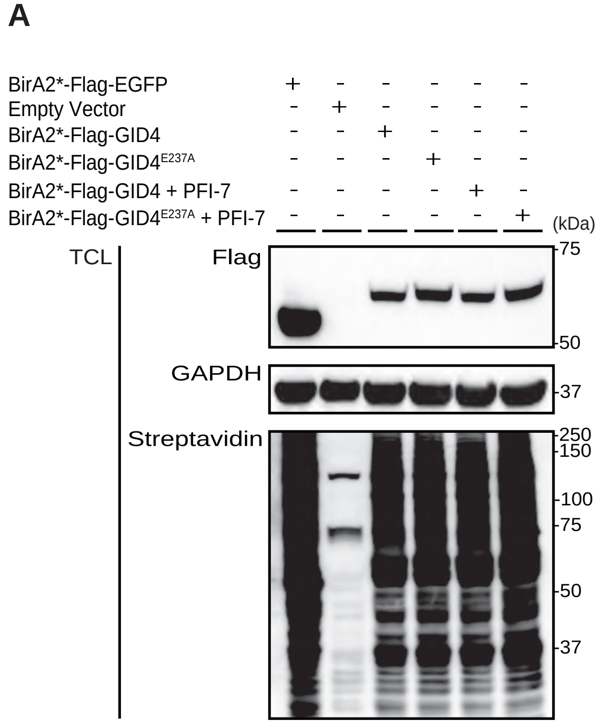

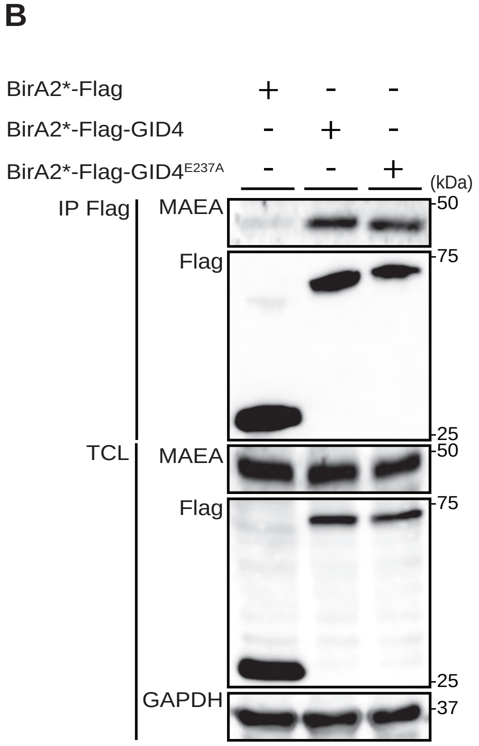

### Figure EV3

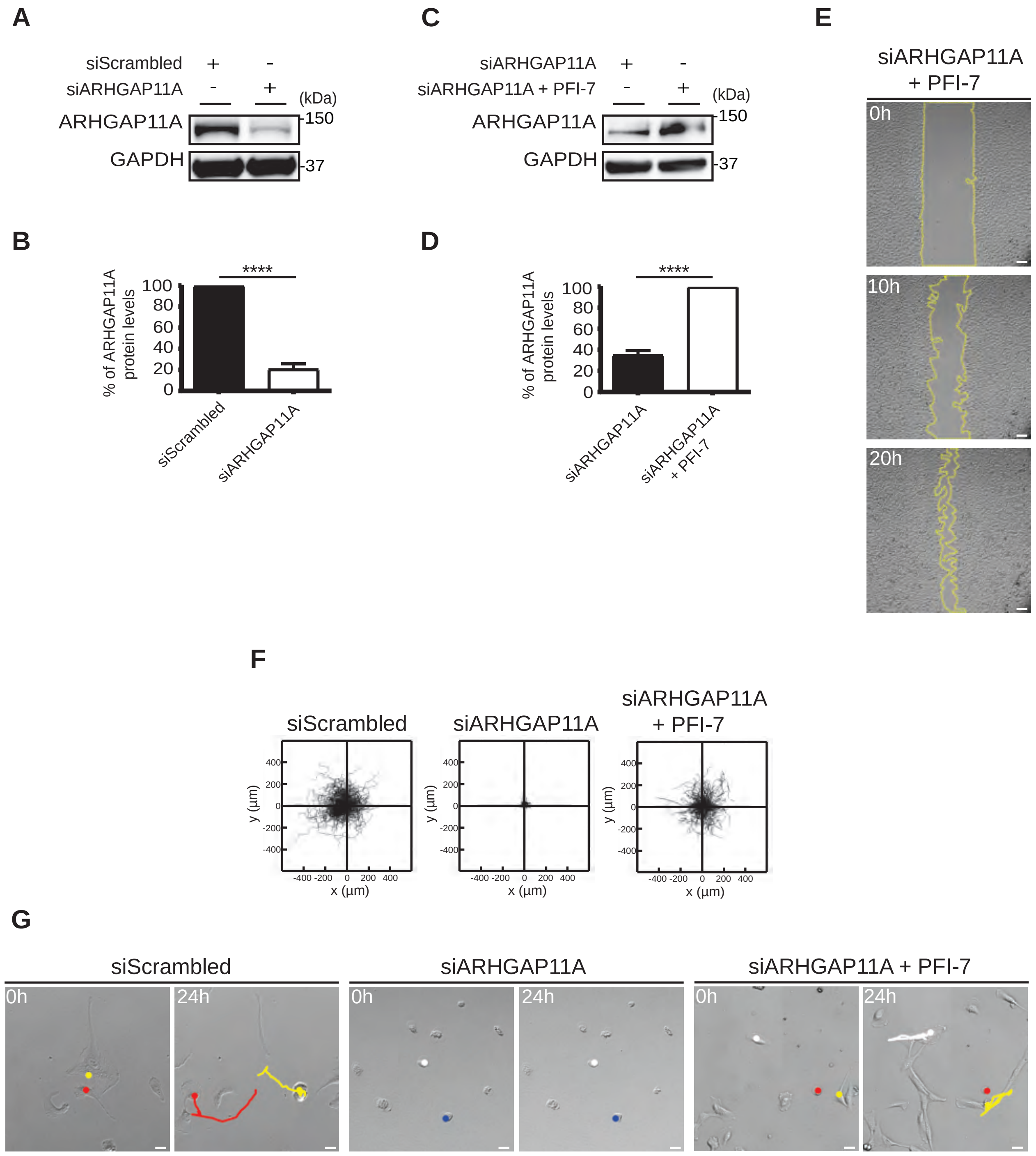

### Figure EV4

#### Fig EV4

Secondary antibody

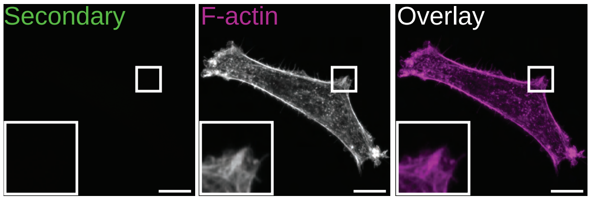
